# Efficacy and mechanism of action of cipargamin as an antibabesial drug candidate

**DOI:** 10.1101/2024.07.15.603500

**Authors:** Hang Li, Shengwei Ji, Nanang R. Ariefta, Eloiza May S. Galon, Shimaa AES El-Sayed, Thom Do, Lijun Jia, Miako Sakaguchi, Masahito Asada, Yoshifumi Nishikawa, Xin Qin, Mingming Liu, Xuenan Xuan

## Abstract

Babesiosis is a disease brought on by intraerythrocytic parasites of the genus *Babesia*. Current chemotherapies are accompanied by side effects and parasite relapse. Therefore, it is crucial to develop highly effective drugs against *Babesia*. Cipargamin (CIP) has shown inhibition against apicomplexan parasites, mainly *Plasmodium* and *Toxoplasma*. This study evaluated the growth-inhibiting properties of CIP against *Babesia* spp. and investigated the mechanism of CIP on *B. gibsoni*. The half inhibitory concentration (IC_50_) values of CIP against the *in vitro* growth of *B. bovis* and *B. gibsoni* were 20.2 ± 1.4 nM and 69.4 ± 2.2 nM, respectively. CIP significantly inhibited the growth of *B. microti* and *B. rodhaini in vivo.* Resistance was conferred by L921V and L921I mutations in BgATP4, which reduced the sensitivity to CIP by 6.1- and 12.8-fold. The inhibitory potency of CIP against BgATP4-associated ATPase activity was moderately reduced in mutant strains, with a 1.3-fold and 2.4-fold decrease in BgATP4^L921V^ and BgATP4^L921I^, respectively, compared to that of BgATP4^WT^. An *in silico* investigation revealed reductions in affinity for CIP binding to BgATP4^L921V^ and BgATP4^L921I^ compared to BgATP4^WT^. Resistant strains showed no significant cross-resistance to atovaquone (ATO) or tafenoquine succinate (TQ), with less than a onefold change in IC_50_ values. Combining CIP with TQ effectively eliminated *B. microti* infection in SCID mice with no relapse, and parasite DNA was not detected by qPCR within 90 days post-infection. Our findings reveal the efficacy of CIP as an anti-babesial agent, its limitations as a monotherapy due to resistance development, and the potential of combination therapy with TQ to overcome said resistance and achieve complete parasite clearance.

## Introduction

*Babesia* is an apicomplexan tick-transmitted hemoparasite, which not only impacts the livestock economy but also causes an emerging disease in humans (Jalovecka et al., 2019). *Babesia* is a global pathogen that is more prevalent in certain regions such as Asia, Europe, and North America. However, as climate change brings with it higher temperatures and humidity, ticks and their reservoir hosts are anticipated to expand northward for survival and activity (Gray and Ogden, 2021). The disease is known as babesiosis, commonly characterized by fever and hemolytic anemia, but chronic infections can be asymptomatic (Almazán et al., 2022). The fatality rate ranges from 1% among all cases to 3% among hospitalized cases, and as high as 20% in immunocompromised patients (Krause, 2019).

Currently, the most common drugs for the treatment of human babesiosis include a combination of atovaquone (ATO) plus azithromycin (AZI) or clindamycin (CLN) plus quinine (QUI) (Krause et al., 2021). Still, these recommended treatments often result in various problems. For instance, multiple mutations in the *B. microti* cytochrome b, which is targeted by ATO, were identified in patients with relapsing babesiosis (Holbrook et al., 2023; Krause et al., 2024; Lemieux et al., 2016; Marcos et al., 2022; Rogers et al., 2023; Rosenblatt et al., 2021; Simon et al., 2017). The combination of CLN and QUI is the last resort for patients with severe symptoms (Kletsova et al., 2017). Despite its efficacy, this combination can elicit adverse drug reactions (Vannier and Krause, 2012). The 8-aminoquinoline analog tafenoquine (TQ) was found to be effective in curing immunocompromised patients experiencing relapsing babesiosis caused by *B. microti* (Rogers et al., 2023). However, the efficacy of TQ treatment may vary between individuals and cases and its contraindication in patients with glucose-6-phosphate dehydrogenase (G6PD) deficiency limits its clinical use (Chu and Freedman, 2019). Due to the aforementioned issues, it is undoubtedly urgent to search for compounds that treat infections caused by *Babesia* spp. while simultaneously minimizing the detrimental side effects of antibabesial drugs.

Spiroindolone cipargamin (CIP), a promising antimalarial, has been found to effectively suppress the growth of all strains of *Plasmodium falciparum* and *P. vivax* with potency in the low nanomolar ranges (Rosling et al., 2018; Rottmann et al., 2010). The assessment of the drug indicated that CIP taken orally had good absorption, a long half-life, and exceptional bioavailability (Schmitt et al., 2022). Following oral CIP treatment (30 mg daily for three days) in adults with simple *P. falciparum* or *P. vivax* malaria, parasitemia was rapidly cleared in a phase II trial (Schmitt et al., 2022). Currently, a clinical trial is being conducted for the intravenous administration of CIP as a treatment for severe malaria patients (ClinicalTrials, 2024). CIP was also evaluated for treating toxoplasmosis as a second-line medicine in cases where there are intolerable toxicity issues or allergies to the currently used treatments. Mice infected with *Toxoplasma gondii* that were treated with CIP on the day of infection and the following day had 90% fewer parasites five days post-infection (Zhou et al., 2014).

*Pf*ATP4, a P-type ATPase in *P. falciparum*, functions as a Na⁺/H⁺ transporter critical for maintaining ionic balance (Spillman et al., 2013). Mutations in *Pf*ATP4 confer resistance to CIP, which disrupts Na⁺ homeostasis, leading to increased cytosolic Na⁺ concentration and various physiological changes, such as cytosolic alkalinization, osmotic swelling, etc. (Mohring et al., 2022). These mutations reduce the Na⁺-dysregulating effects of CIP and the resting Na⁺ levels. While *Pf*ATP4 is essential for *P. falciparum* survival, its homolog in *T. gondii* (*Tg*ATP4) is less critical, as *T. gondii* experiences brief, high Na⁺ exposure and can survive without *Tg*ATP4 expression (Lehane et al., 2019).

Due to its excellent efficacy against other apicomplexan parasites, including *Plasmodium* spp., and its ability to target parasite ATP4—a conserved protein essential for ion homeostasis in apicomplexan organisms—we hypothesize that CIP could also effectively inhibit *Babesia* spp., a related apicomplexan parasite. Therefore, the objectives of this study were to determine whether CIP could inhibit the growth of *Babesia* spp., namely *B. bovis* and *B. gibsoni in vitro* and *B. microti* and *B. rodhaini in vivo*, and identify the inhibitory mechanisms of CIP on *Babesia* parasites.

## Results

### Inhibitory efficacy of CIP on *B. bovis* and *B. gibsoni in vitro*

*In vitro* efficacy of CIP against *B. bovis* and *B. gibsoni* showed a steep growth inhibition curve with half inhibitory concentration (IC_50_) values of 20.2 ± 1.4 nM (Figure 1A) and 69.4 ± 2.2 nM (Figure 1B), respectively. The 50% cytotoxic concentration (CC_50_) value of CIP on Madin-Darby canine kidney (MDCK) cells and human foreskin fibroblasts (HFF) were 38.7 ± 2.0 μM and 70.8 ± 4.9 μM (Figure supplement 1), respectively. Based on these values, the predicted selectivity indices (SIs), which reflect the drug’s safety and specificity, were calculated to be greater than 500. Furthermore, at a concentration of 100 μM, CIP exhibited a low erythrocyte hemolysis rate of 0.11 ± 0.03 % (data not shown).

**Figure 1.**
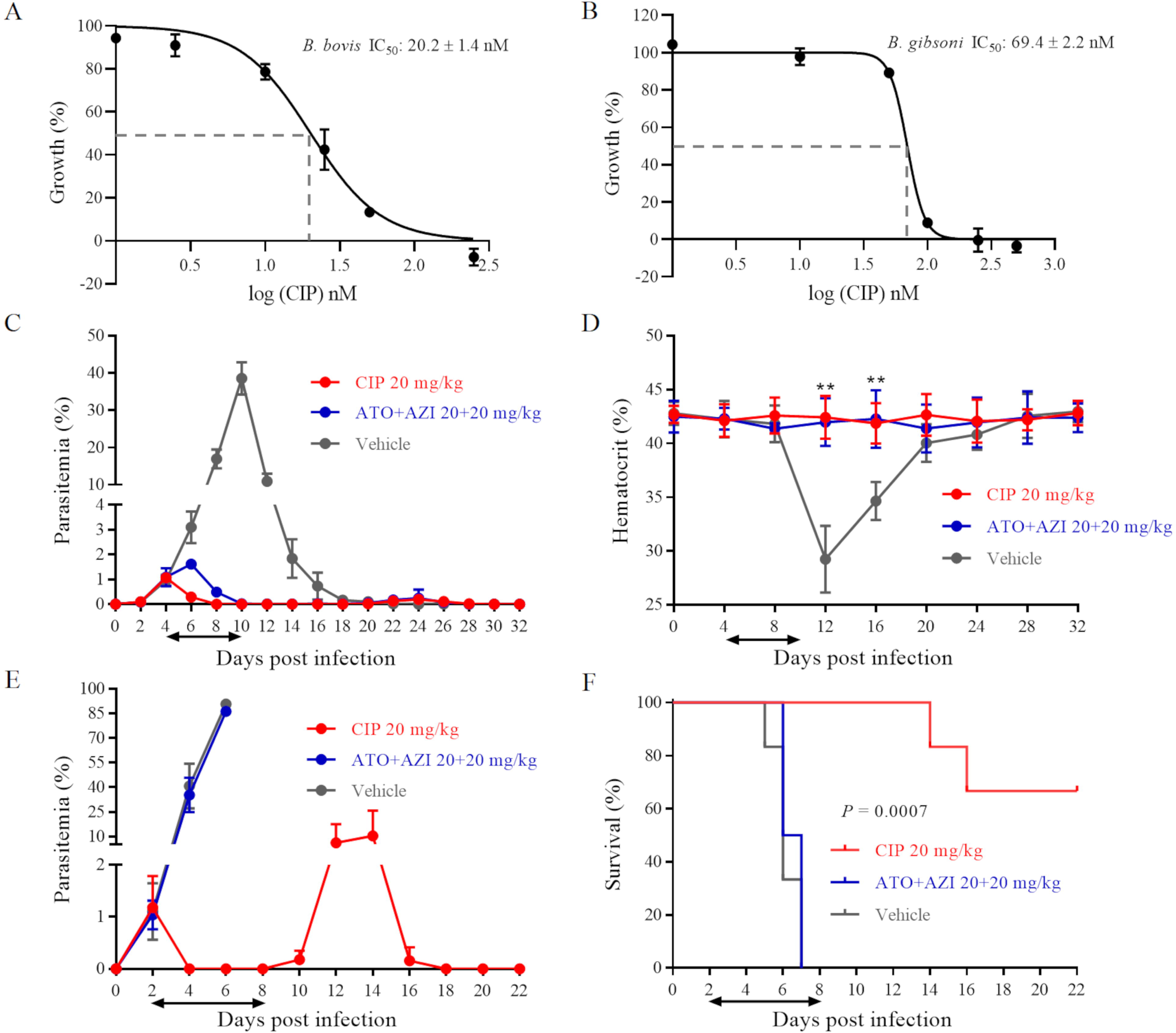
CIP demonstrates potent inhibition on *Babesia* spp. **(A)** and (**B)** Dose-dependent growth curve of CIP on *B. bovis* and *B. gibsoni in vitro*. Each value represents the mean ± standard deviation (SD) of three independent experiments carried out in triplicate. IC_50_: the half maximal inhibitory concentration. (**C)** Inhibitory effects of CIP and atovaquone (ATO) plus azithromycin (AZI) on the proliferation of *B. microti* in BALB/c mice. (**D)** Hematocrit (HCT) values in mice treated with CIP or ATO plus AZI compared with vehicle-treated mice. (**E)** Inhibitory effects of CIP and ATO plus AZI on the proliferation of *B. rodhaini* in BALB/c mice. (**F)** Survival rates of CIP-treated, ATO plus AZI-treated, and vehicle-treated mice. The treatment time is shown by two-way arrows, and significant differences (*P* < 0.01) between the drug-treated groups and the vehicle-treated control group are indicated by asterisks. The data from one of six individual experiments are expressed as means ± SD. **, *P* ˂ 0.01.

### CIP effect on B. microti and B. rodhaini infections in vivo

Concurrently, CIP showed effective inhibition on *B. microti* and *B. rodhaini in vivo*. The parasitemia of *B. microti*-infected BALB/c mice increased dramatically in the vehicle-treated control group and peaked at 10 days postinfection (DPI) (38.55 ± 4.32%) (Figure 1C). On the other hand, seven days of treatment with CIP (20 mg/kg) or atovaquone (ATO) plus azithromycin (AZI) administered orally resulted in a significantly lower peak parasitemia, 1.06 ± 0.20% and 1.61 ± 0.20%, respectively (Figure 1C). Hematocrit (HCT) variations were tracked every 4 days as an indicator of anemia in *B. microti*-infected mice. The vehicle-treated group showed a drop in HCT levels at 12 DPI and 16 DPI (*P* < 0.01) (Figure 1D). No significant reduction in HCT levels was observed in the CIP-treated group or the ATO plus AZI-treated group (Figure 1D). This indicates that the administration of CIP could control *B. microti* infection and prevent anemia from developing in *B. microti*-infected mice. BALB/c mice infected with *B. rodhaini* treated with sesame oil or ATO plus AZI showed high parasitemia, 90.73 ± 1.97%, and 86.23 ± 3.06%, respectively (Figure 1E), and all mice died within 7 DPI (Figure 1F). CIP treatment in *B. rodhaini*-infected mice precluded the emergence of parasitemia for the following eight days (Figure 1E), which led to 66.67% of mice surviving the challenge infection (Figure 1F). At 12 DPI, parasites had recurred in all CIP-treated *B. rodhaini*-infected mice (10.32 ± 15.51%), which were eventually cleared as indicated by undetectable parasites at 18 DPI (Figure 1E).

### Identification of *B. gibsoni* ATP4 mutation in CIP-resistant strains

After being exposed to CIP at increasing concentrations up to 10 times the IC_50_, the resistant parasites in two of the culture wells were able to regrow. We sequenced the *B. gibsoni* ATP4 gene from the wild-type and two resistant strains. The wild-type strain has a C at nucleotide 2,761, which translates to leucine (Figure 2A). In one resistant strain, a single nucleotide variant (SNV) in BgATP4 with a substitution at position 2,761 (from C to G) was found − a nonsynonymous coding change from leucine to valine (L921V) (Figure 2B). In another resistant strain, the mutation occurred in the same position. However, the nucleotide substitution was from C to A, and the coding changed from leucine to isoleucine (L921I) (Figure 2C). Next-generation sequencing (NGS) revealed that for *Bgatp4*^2761C>G^, 99.97% of 7,960 reads were G at nucleotide 2,761, and for *Bgatp4*^2761C>A^, 99.92% of 7,862 reads were A at nucleotide 2,761 (Figure 2D). BgATP4^L921V^ and BgATP4^L921I^ lines were tested for their susceptibility to CIP, and had IC_50_ values of 421.0 ± 15.9 nM and 887.9 ± 62.0 nM, respectively (Figure 2E). These findings demonstrate a 6.1- and 12.8-fold reduction in CIP sensitivity of the resistant parasite lines BgATP4^L921V^ and BgATP4^L921I^.

**Figure 2.**
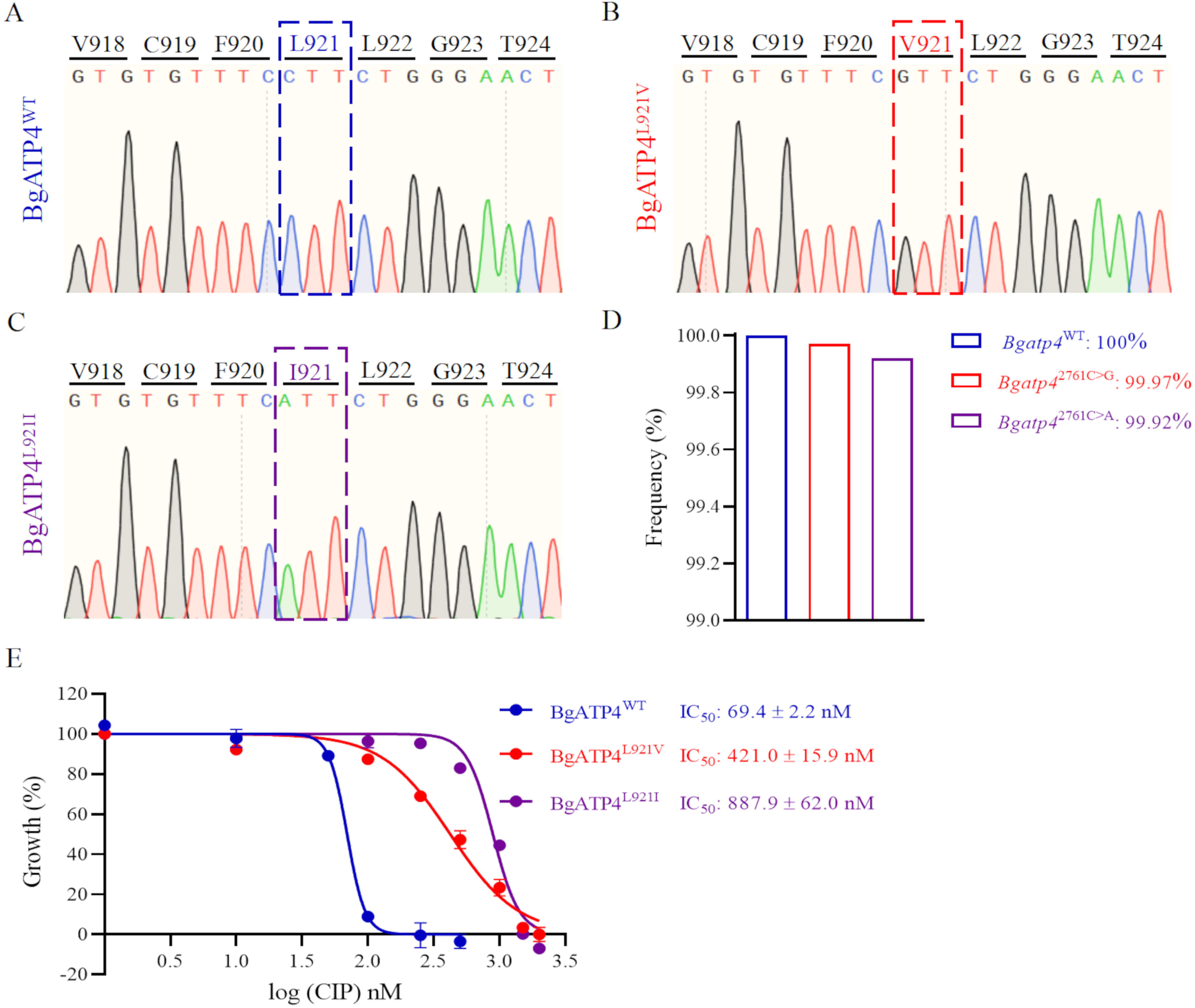
Mutations in BgATP4 mediate CIP resistance. **(A**-**C)** Representative sequencing chromatogram of wild-type and resistant parasites from CIP-treated *B. gibsoni*. The resistant parasite genomic DNA is extracted from blood samples after a 60 day-treatment. The BgATP4 gene was amplified and sequenced using the DNA. (**D)** Genes of high-frequency sequence variants detected by NGS. (**E)** Dose-dependent growth curve of BgATP4^WT^, BgATP4^L921V^, and BgATP4^L921I^ *in vitro*. Each value represents the mean ± standard deviation (SD) of three independent experiments carried out in triplicate.

### The effect of CIP on BgATP4^WT^, BgATP4^L921V^ and BgATP4^L921I^ function

Microscopic observation of thin blood smears was performed to determine the morphological changes of *B. gibsoni* exposed to CIP. The CIP-treated parasites became swollen after incubation with the drug for 72 h (Figure 3A). For both the treatment and control groups, one hundred parasites were measured. The mean size of treated parasites was notably bigger than the parasites in the untreated group (*P* ˂ 0.0001) (Figure 3B). Significant vacuolization was observed in the cytoplasm of parasites in the CIP-treated group, as revealed by transmission electron microscopy (TEM). Despite this, the nuclear membrane structure and parasitic membranes remained intact until the parasites were completely destroyed (Figure 3C). The addition of the ATP4 inhibitor CIP resulted in a time-dependent increase in the concentrations of [Na^+^]_i_ in wild-type *B. gibsoni*, with improved signal-to-noise ratios at the higher drug concentration of 20 nM (Figure 3D). We observed that the Na^+^ concentrations in both BgATP4^L921V^ and BgATP4^L921I^ lines were lower when compared with those of the control BgATP4^WT^ line after being exposed to 20 nM CIP for 20 min, with a significantly lower Na^+^ concentration in BgATP4^L921I^ (*P* = 0.0087) (Figure 3F). We also demonstrated here that the addition of CIP in wild-type *B. gibsoni* caused an increase in the cytosolic pH (Figure 3E). Specifically, 4 min after the drug was added, the average pH of the 20 nM CIP group reached as high as 7.278, while the 1 nM CIP group reached 7.089 and the untreated group reached 7.062 (Figure 3E). The pH values of the 20 nM CIP group were consistently higher than those of the other two groups, although declining with time (Figure 3E). In resistant lines, a 20-min exposure to 20 nM CIP caused small changes in the pH values compared to the wild-type line, with the BgATP4^L921I^ line (7.048 ± 0.042) having a notably lower pH value (*P* = 0.0229) (Figure 3G).

**Figure 3.**
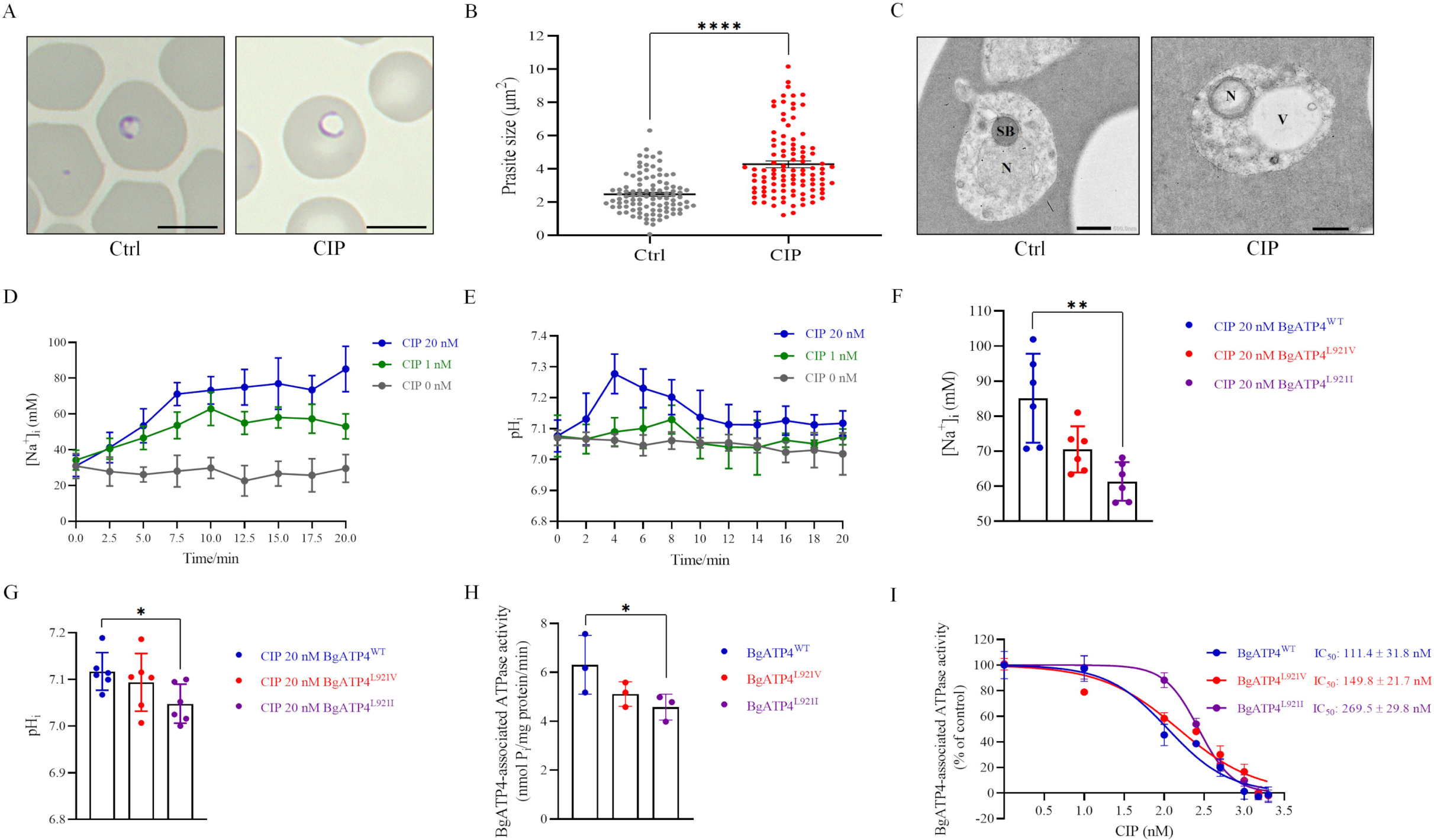
Mechanistic basis for resistance to CIP conferred by the L921V and L921I mutations in BgATP4. **(A)** Untreated and CIP-treated parasite morphology after incubation for 72 h. Scale bar: 5 µm. (**B)** Sizes of 100 parasites in two groups measured with ImageJ software in panels A. Statistically significant differences between the means of variables determined by t-test. ****, *P* ˂ 0.0001. (**C)** TEM of untreated and CIP-treated parasite. N, Nucleus; SB, spherical body; V, vacuole. Scale bar: 500 nm. (**D)** [Na^+^]_i_ concentrations after the addition of CIP in the BgATP4^WT^ line. Representative traces from the experiment that highlight the impact of adding 20 nM CIP (blue), 1 nM CIP (green), or 0 nM CIP (grey) on the concentration [Na^+^]_i_ of the BgATP4^WT^ line. (**E)** Alkalinization of pH_i_ in BgATP4^WT^ line upon addition of the ATP4 inhibitor. (**F)** Addition of 20 nM CIP to the wild-type and resistant parasite lines results in different [Na^+^]_i_ concentrations. (**G)** Addition of 20 nM CIP to the wild-type and resistant parasite lines results in different pH_i_ concentrations. Experiments were performed in technical duplicates for at least three biological repeats. **(H)** Data acquired in the low Na^+^ condition (containing only the 2 mM Na^+^ introduced upon the addition of 1 mM Na_2_ATP) was subtracted from data obtained in the high Na^+^ condition to determine the ATPase activity related to the BgATP4 proteins. The results are presented as the average of data from three independent tests. (**I)** Dose-dependent BgATP4-associated ATPase activity curve of BgATP4^WT^, BgATP4^L921V^, and BgATP4^L921I^ *in vitro*. ATPase activity was determined at pH 7.2 in the presence of 150 mM Na^+^ and 1 mM Na_2_ATP. Each value represents the mean ± standard deviation (SD) of three independent experiments carried out in triplicate. *, *P* ˂ 0.05; **, *P* ˂ 0.01.

### Sensitivity of BgATP4-associated ATPase activity to CIP in BgATP4^WT^, BgATP4^L921V^, and BgATP4^L921I^

The BgATP4-associated ATPase activity in erythrocytes infected with BgATP4^WT^ (6.31 ± 1.20 nmol Pi/mg protein/min), measured in the presence of 150 mM Na^+^, was higher than those observed in BgATP4^L921V^ (5.11 ± 0.50 nmol Pi/mg protein/min) and BgATP4^L921I^ (4.58 ± 0.53 nmol Pi/mg protein/min) (*P* = 0.04) (Figure 3H).

We further investigated the concentration-dependent inhibition of BgATP4-associated ATPase activity by CIP in wild-type and mutant parasites. In membranes prepared from *B.gibsoni*, CIP inhibited BgATP4-associated ATPase activity with IC_50_ values of 111.4 ± 31.8 nM for BgATP4^WT^, 149.8 ± 21.7 nM for BgATP4^L921V^, and 269.5 ± 29.8 nM for BgATP4^L921I^. The potency of CIP in inhibiting BgATP4-associated ATPase activity was reduced by 1.3-fold and 2.4-fold in membranes prepared from BgATP4^L921V^ and BgATP4^L921I^, respectively, compared to BgATP4^WT^ (Figure 3I).

### Multiple sequence alignment of *Babesia* ATP4 and molecular docking

The whole amino acid sequence of *B. gibsoni* ATP4 (GenBank: KAK1443404.1) shared identity values of 29.75%, 49.40%, 49.67%, 62.21%, and 52.47% with *Homo sapiens* ATP4 (GenBank: NM_000704.3), *P. falciparum* ATP4 (GenBank: PF3D7_1211900), *T. gondii* ATP4 (GenBank: XP_018635122.1), *B. bovis* ATP4 (PiroplasmaDB: BBOV_IV010020), and *B. microti* ATP4 (GenBank: BMR1_03g01005), respectively (Figure supplement 2).

The pLDDT (predicted Local Distance Difference Test) value of BgATP4^WT^ prediction was 80.7 using Colab-fold. Multiple potential binding sites for CIP were revealed by blind docking throughout the whole protein surface (Figure supplement 3). CIP binds in close proximity to L921, as demonstrated by focused docking on this area (Figure 4A). The contribution of each residue to the predicted binding affinity in either mutant structure was reduced; the precise values of BgATP4^WT^ (Figure 4B), BgATP4^L921V^ (Figure 4C), and BgATP4^L921I^ (Figure 4D) were -6.43, -6.40 and -6.26 kcal/mol, respectively. The interactions of CIP from each docking simulation are shown in Table supplement 3.

**Figure 4.**
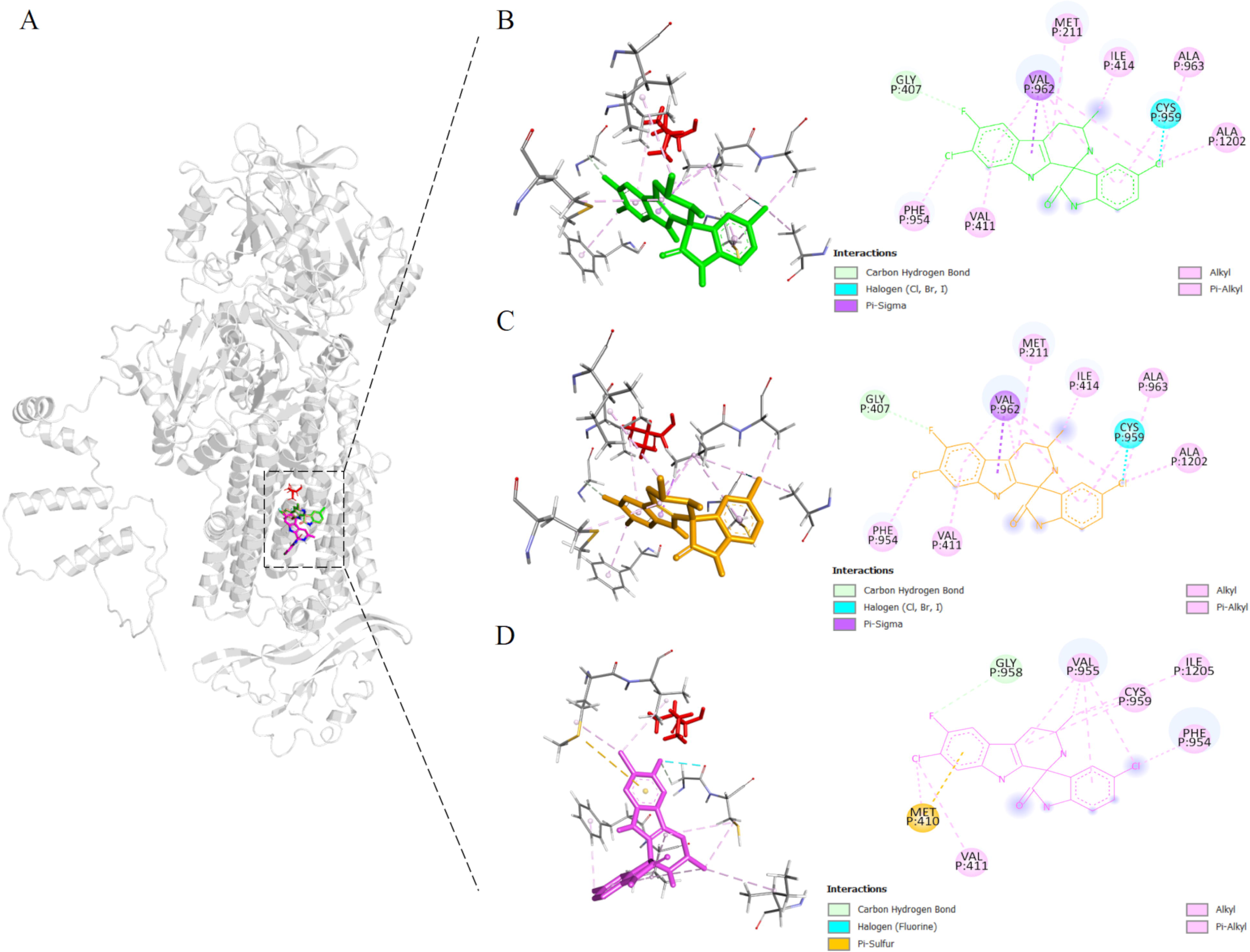
Binding sites proximal to BgATP4 residue 921 predicted by molecular docking. **(A)** The lowest energy poses for CIP were located in reference to the whole protein structure, docking against the WT (green), L921V (yellow), and L921I (pink) mutant BgATP4. The side chain of L921 is also shown in a red stick at its position. (**B**-**D)** The zoomed views of the binding locations of CIP.

### Cross-resistance of BgATP4^L921V^ and BgATP4^L921I^ mutants to ATO and TQ

The BgATP4^WT^, BgATP4^L921V^, and BgATP4^L921I^ lines were tested for their susceptibility to ATO and TQ. The IC_50_ values for these lines were 380.1 ± 3.1 nM, 412.9 ± 5.4 nM, and 360.3 ± 7.9 nM, respectively, for ATO (Figure 5A), and 39.5 ± 0.4 μM, 29.9 ± 1.4 μM, and 39.3 ± 0.3 μM, respectively, for TQ (Figure 5B). The IC_50_ values for both ATO and TQ in the resistant strains showed only slight changes compared to the wild-type strain, with less than a onefold difference. This minimal variation suggests that the resistant strain has a mild alteration in susceptibility to ATO and TQ, but not enough to indicate strong resistance or significant cross-resistance.

**Figure 5.**
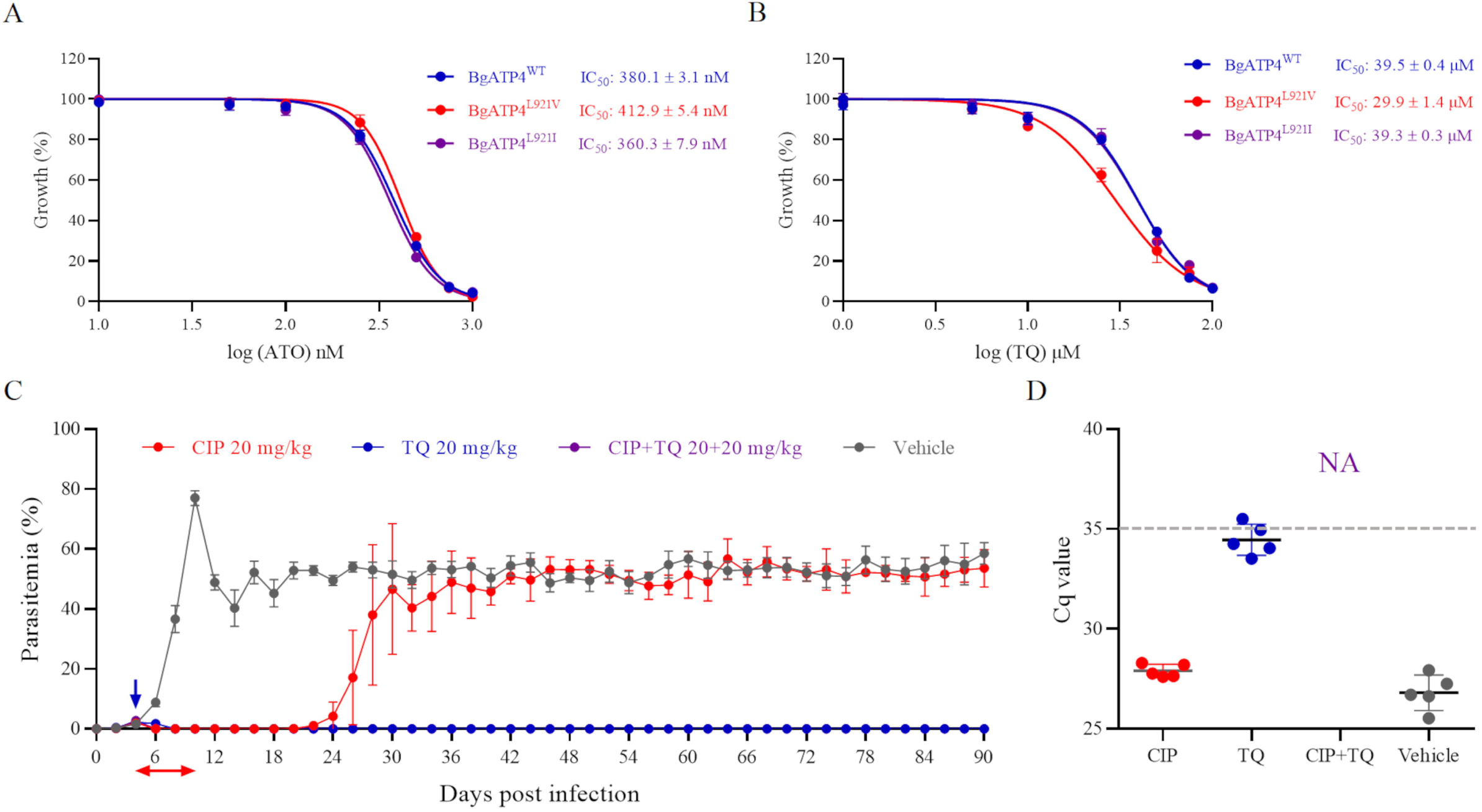
Cross-resistance between ATO and TQ in resistant parasites and combination therapy based on CIP plus TQ. **(A)** Dose-dependent growth curve of BgATP4^WT^, BgATP4^L921V^, and BgATP4^L921I^ by ATO treatment *in vitro*. **(B)** Dose-dependent growth curve of BgATP4^WT^, BgATP4^L921V^, and BgATP4^L921I^ by TQ treatment *in vitro.* Each value represents the mean ± standard deviation (SD) of three independent experiments carried out in triplicate. **(C)** Inhibitory effects of CIP plus TQ on the proliferation of *B. microti* in SCID mice. Treatment started at 4 DPI: CIP was given at 20 mg/kg once daily for 7 days, TQ was administered as a single 20 mg/kg dose, and the combination group received both treatments (CIP at 20 mg/kg once daily for 7 days plus a single dose of TQ at 20 mg/kg). The data are presented as the means from one of five independent experiments. **(D)** Parasite DNA was detected by qPCR on genomic DNA extracted from blood collected from untreated and treated SCID mice infected with *B. microti* at 90 DPI. A dotted grey line across the graph represents the average cut-off Cq value. Cut-off Cq ≤ 35 was considered positive, while Cq > 35 or no amplification was considered negative. Each sample was analyzed in duplicate, and two independent experiments were conducted. NA indicates no amplification.

### Combination treatment of CIP plus TQ in SCID mice with *B. microti* infection

The parasitemia of *B. microti*-infected SCID mice in the vehicle group increased dramatically, peaking at 10 DPI (77.03 ± 2.45%) (Figure 5C). Although the trend showed a subsequent decline, the parasitemia remained around 50% until the mice were euthanized at 90 DPI. Treatment with CIP (20 mg/kg) initially reduced parasitemia to undetectable by 18 DPI. However, a relapse occurred, with parasitemia increasing rapidly and stabilizing at levels comparable to the vehicle group during the subsequent observation period. Notably, no mutations were detected in the relapsed *B. microti* parasites from SCID mice (data not shown). In contrast, parasitemia was completely cleared in the TQ and CIP plus TQ groups by 8 DPI and 6 DPI, respectively, with no parasites observed under the microscope thereafter. qPCR analysis at 90 DPI detected parasite DNA in all groups except the CIP plus TQ group (Figure 5D).

## Discussion

The repositioning of antimalarial drugs is critical in developing novel strategies for treating babesiosis. CIP is a novel compound that inhibits *Plasmodium* development by targeting ATP4 and has been extensively tested in Phase 1 and Phase 2 clinical trials (Qiu et al., 2022). Since ATP4 is conserved across apicomplexans, including *Babesia* species, our study aimed to repurpose the antimalarial CIP by assessing its efficacy on *Babesia* species (Lehane et al., 2019). The IC_50_ values of CIP against *B. bovis* and *B. gibsoni in vitro* were lower than the previously reported IC_50_ value of TQ (Carvalho et al., 2020; Ji et al., 2022b) and ATO against *B. gibsoni* (Matsuu et al., 2004). The present investigation also demonstrated that the inhibitory effects of CIP on *B. microti*-infected BALB/c mice were comparable to that of ATO plus AZI, the combination recommended by the CDC in the United States. CIP was also proven to protect mice from the deadly *B. rodhaini* infection, with a survival rate of up to 67%, as recorded in the current trial. These results suggest that CIP may be a potential chemotherapy candidate for babesiosis.

In a previous study, the *P. falciparum* Dd2 strain that acquired resistance to CIP carried the G358S mutation in the *Pf*ATP4 protein, a *P. falciparum* P-type Na^+^ ATPase (Qiu et al., 2022). The vital protein ATP4 is found in the parasite plasma membrane and is specific to the subclass of apicomplexan parasites (Mohring et al., 2022). To date, *in vitro* evolution experiments using CIP have produced at least 18 parasite lines with various *Pf*ATP4 mutations (Lee and Fidock, 2016; Rottmann et al., 2010). In another study involving *T. gondii*, a cell line that carried the mutation G419S in the *Tg*ATP4 gene was 34 times less susceptible to CIP than that of *Tg*ATP4^WT^ (Qiu et al., 2022). It follows that there is a significant possibility that resistant *Babesia* parasites will emerge from CIP exposure. In this study, we successfully produced two CIP-resistant strains using six independent selections. The newly discovered L921V and L921I mutations in BgATP4 decreased the CIP sensitivity by 6.1 and 12.8 times, respectively. We provided compelling evidence that natural variety mutations L921V and L921I in BgATP4 significantly affected the protein’s susceptibility to CIP inhibition, albeit the mutational sites were different from those of *P. falciparum* and *T. gondii*. Based on our observation, the growth and generation rates of the mutant strains are comparable to those of the wild-type strain.

Although ATP4 was initially recognized as a Ca^2+^ transporter (Krishna et al., 2001), current evidence suggests that ATP4 functions as an ATPase for exporting Na^+^ while importing H^+^ (Mohring et al., 2022), as well as causing a variety of other physiological perturbations including an increase in the volume of parasites and infected erythrocytes due to the osmotic impact of the [Na^+^]_cyt_ increase (Dennis et al., 2018), a decline in cholesterol extrusion from the parasite plasma membrane as a result of the increase in [Na^+^]_cyt_ (Das et al., 2016), and an intensified rigidity of erythrocytes infected with ring-stage parasites (Zhang et al., 2016). In this study, the CIP-exposed wild-type *B. gibsoni* became swollen, which was identical to a prior study on *P. falciparum* (Dennis et al., 2018). Interestingly, transmission electron microscopy (TEM) analysis of parasites incubated with CIP revealed ultrastructural alterations characterized by significant vacuole formation in the cytoplasm, a hallmark of stress or early stages of cell death. These changes were similar to those observed in *B. bovis* treated with clotrimazole and ketoconazole, which ultimately led to growth inhibition or parasite death (Bork et al., 2003). Despite the vacuolization, the structural integrity of the nucleus and membranes was maintained, suggesting a gradual process where vacuolization precedes complete destruction of the parasite. One explanation is the swelling of the isolated parasites, which can be ascribed to the osmotic consequences of Na^+^ uptake and is contingent upon the presence of Na^+^ in the external environment.

The output of Na^+^ and input of H^+^ diminished upon CIP-induced inhibition of *Pf*ATP4, and the continued outflow of H^+^ via V-type H^+^-ATPase led to an alkalinization that ultimately killed the parasites (Spillman et al., 2013). Our capacity to measure a time-dependent increase in the concentration of [Na^+^]_i_ and pH value for wild-type *B. gibsoni* facilitated us to gain insight into the basic mechanisms of CIP on BgATP4^WT^ function. Furthermore, our results corroborate internal alkalinization as the main factor in *Babesia* death and support our hypothesis that the swollen isolated parasites were produced by Na^+^ absorption. To explore further how mutations in BgATP4 are associated with the upregulation of the parasite’s [Na^+^]_i_ and [H^+^]_i_, we tested two BgATP4-mutant lines that were chosen previously with BgATP4 inhibitors (BgATP4^L921V^ and BgATP4^L921I^). Due to the presence of L921V and L921I mutations in drug-resistant strains of BgATP4, the concentration of Na^+^ did not increase as much as it would have in the wild-type strain following the addition of 20 nM CIP for 20 min. As intraerythrocytic alkalinization rises, the same outcome happens. From these results, we deduced how natural mutations in BgATP4 may affect ATP4 inhibitor susceptibility by dysregulating H^+^ and Na^+^ balance, which helped parasites survive in a relatively high concentration of CIP. An illustration of the putative processes for regulating Na^+^ and H^+^ in erythrocytes infected with the *Babesia* parasite is presented in Figure supplement 4.

In a previous study’s isolated membrane assay, CIP was found to be more effective at inhibiting parasite development than *Pf*ATP4-associated ATPase activity and reduced sensitivity of *Pf*ATP4-associated ATPase activity to the medication was associated with a decrease in parasite susceptibility to CIP (Rosling et al., 2018). In our study, the higher resistance to CIP-mediated inhibition of parasite proliferation observed in BgATP4^L921I^ compared to BgATP4^L921V^ correlated with a higher resistance of BgATP4-associated ATPase activity to the drug. These findings provide additional evidence that the ATPase activity measured in this study corresponds to the activity of the BgATP4 protein.

According to studies using the *Pf*ATP4 model for molecular docking, the G358S mutation results in a steric clash that lowers CIP’s binding affinity (Qiu et al., 2022). The results from the current study were similar to the *Pf*ATP4 model. The molecular docking was constructed using a ColabFold model of wild-type BgATP4, which predicted the binding mode and affinity between ATP4 protein and the ligand to provide a possible mechanistic explanation. It suggested that the L921V mutation caused changes at the atomic level, whereas the L921I mutation created a steric clash that reduced the binding affinity of CIP. The predicted affinity score of the L921I mutation was lower than that of the L921V mutation. Thus, it is possible that CIP had a weaker binding to BgATP4^L921I^ than to BgATP4^L921V^. These findings are consistent with the results obtained from measuring the IC_50_ of the drug against parasites with the L921V and L921I mutations.

While CIP shows promise as an antibabesial agent, the emergence of resistance emphasizes the importance of rational drug use and the development of combination therapies. Our results demonstrate that CIP did not exhibit cross-resistance with ATO or TQ, as IC_50_ values for ATO and TQ in resistant strains showed minimal changes (<1-fold) compared to the wild-type strain. This observation suggests that CIP could be effectively combined with other antibabesial drugs to reduce the risk of resistance development. Previous studies have also reported that a single dose of TQ may not be sufficient to prevent parasite relapse in immunocompromised hosts, recommending either repeated TQ administration or combination therapy with other antibabesial agents to address this issue (Mordue et al., 2019). Supporting this approach, our study demonstrates that the combination of CIP plus TQ effectively cleared *B. microti* infections in SCID mice, with no detectable parasitemia or DNA observed at 90 DPI, indicating complete parasite elimination.

This preclinical finding underscores the potential of the CIP plus TQ combination as a viable treatment strategy for babesiosis, especially in cases where monotherapy is ineffective. Further studies are needed to fully explore the pharmacokinetics, long-term efficacy, and potential side effects of such combinations in clinical settings, ultimately facilitating optimized therapeutic strategies.

In summary, our findings provide a comprehensive understanding of CIP’s efficacy, mechanism of resistance, and identifying strategies to enhance its therapeutic potential. The combination of CIP with drugs like TQ offers a promising approach to combat babesiosis and mitigate the development of resistance.

## Materials and methods

### Parasite culture

The parasites *B. gibsoni* Oita strain and *B. bovis* Texas strain were *in vitro* cultured in 24-well plates and maintained in an atmosphere of 5% CO_2_ and 5% O_2_ at 37 °C (Liu et al., 2018). For the *in vivo* studies, *B. microti* Peabody mjr strain-(ATCC PRA-99^TM^) and *B. rodhaini* Australia strain-infected RBCs (iRBCs), which were collected and diluted with phosphate-buffered saline (PBS) when the parasitemia levels in the donor mice reached ∼20% and 50%, respectively, and were intraperitoneally injected into BALB/c mice. Each BALB/c mouse was infected with 1.0 × 10^7^ *B. microti* or *B. rodhaini* iRBCs for the *in vivo* trials (Ji et al., 2022a).

### *In vitro* cytotoxicity of CIP and hemolysis rate in canine erythrocytes

Cultures of Madin-Darby canine kidney (MDCK) cells and human foreskin fibroblasts (HFF) were maintained at 37 °C under an atmosphere of 5% CO_2_ and 5% O_2_ and the cytotoxic effect of CIP (MedChem Express, Tokyo, Japan) was assessed using a cell viability assay by CCK-8 (Dojindo, Japan) as described previously (Li et al., 2023). The selectivity index is calculated as the ratio between the half maximal inhibitory concentration (IC_50_) and the 50% cytotoxic concentration (CC_50_) values.

Canine erythrocytes were collected from healthy beagle dogs raised in NRCPD and stocked in Vega y Martinez (VYM) phosphate-buffered saline solution at 4 °C (Vega et al., 1985). A canine erythrocyte hemolysis assay was performed at concentrations of 0.1, 1, 5, 10, 25, 50, and 100 µM as previously described (Ariefta et al., 2022).

### Evaluation of the efficacy of CIP against *Babesia* parasites *in vitro*

The efficacy of CIP against *B. gibsoni* and *B. bovis* was determined using a fluorescence assay, as previously described (Guswanto et al., 2014). The IC_50_ values were determined from the fluorescence values and by non-linear regression analysis (curve fit) in GraphPad Prism 9 (GraphPad Software Inc., USA).

### Chemotherapeutic effects of CIP against *Babesia* infections *in vivo*

CIP was evaluated on *B. microti*- and *B. rodhaini*-infected mice, as previously described (Ndayisaba et al., 2021; Tuvshintulga et al., 2022). When *B. microti*- and *B. rodhaini*-infected mice had a 1% average parasitemia at 4 and 2 days postinfection (DPI), respectively, the drug treatments were administered and continued for seven days. Three groups of *B. microti*-infected mice were administered different treatments. The CIP group (n = 6) and the ATO plus AZI group (n = 6) were orally treated with 20 mg/kg CIP and 20 mg/kg ATO plus 20 mg/kg AZI (Sigma, Tokyo, Japan), respectively. All drugs were prepared in sesame oil. The infected mice of the vehicle group (n = 6) orally received 0.2 mL of sesame oil as the control. Eighteen mice infected with *B. rodhaini* were likewise placed into three groups for treatment, and they received the same treatments as the mice infected with *B. microti*. A light microscope (Nikon, Japan) and a hematology analyzer (Celltac α MEK-6450, Nihon Kohden Corporation, Tokyo, Japan) were used to assess the parasitemia and hematocrit levels every two and four days, respectively.

### Selection of CIP-resistant *B. gibsoni in vitro*

Selections were initiated by exposing six independent flasks, each containing 10 μL (5×10^6^) *B. gibsoni* iRBCs mixed with 40 μL (4×10^8^) RBCs into a 450 μL culture medium, which contained increasing concentrations of CIP: 5, 10, 20, 30 to 694 nM (10×IC_50_). The medium containing CIP was replaced daily until parasites treated with 10×IC_50_ CIP reached multiplication rates that were approximately comparable to those of the untreated controls (Hwang et al., 2010). Then, to evaluate the decreased sensitivity to CIP, IC_50_ values of resistant strains were determined by nonlinear regression using the GraphPad Prism software.

### Detection of *B. gibsoni* ATP4 gene mutations

The genomic DNA of the mutant parasites was extracted and sequenced (Ji et al., 2022a). The primer sets used for sequencing are listed in Table supplement 1. The single nucleotide variants were identified by pairwise alignment to the BgATP4^WT^ sequence (GenBank: JAVEPI010000002.1). Next-generation sequencing (NGS) was performed to detect the ratio of wild-type to mutant parasites by using the Illumina NovaSeq6000 sequencing platform (Jeon et al., 2021).

### Morphological changes in CIP-treated *in vitro* cultured *B. gibsoni*

A microscopy assay was used to detect the morphological changes of wild-type *B. gibsoni* after exposure to 50 nM CIP for three consecutive days (Liu et al., 2021; Tayebwa et al., 2018). At 4 days post-treatment, ImageJ software was used to measure the sizes of 100 randomly selected parasites in the CIP-treated group and the control group on Giemsa-stained blood smears. After treating the parasites as described above, iRBCs were fixed in GA fixation buffer (2% glutaraldehyde in 0.1 M sodium cacodylate buffer containing 1 mM CaCl_2_ and 1 mM MgCl_2_), then stored in rinse buffer (0.1 M sodium cacodylate buffer containing 1 mM CaCl_2_ and 1 mM MgCl_2_) at 4 °C for transmission electron microscopy (TEM) analysis (Hidayati et al., 2023).

### Parasites [Na^+^]_i_ and pH_i_ measurements

For both [Na^+^]_i_ and pH_i_ measurements, the wild-type and mutant parasites were initially separated from erythrocytes by treatment with saponin (0.05% [wt/vol]) for 5 min (Saliba and Kirk, 1999). The Na^+^-sensitive fluorescent dye SBFI (Thermo Fisher Scientific; product S1263) was used to quantify intracellular sodium ([Na^+^]_i_). Saponin-isolated parasites (at 1×10^8^ parasites/mL) were loaded with SBFI (5 μM; in the presence of 0.02% w/v Pluronic F127) for 1 hr at room temperature (RT) (Spillman et al., 2013). Thereafter, SBFI-loaded parasites were resuspended in physiological saline (120 mM NaCl, 5 mM KCl, 25 mM HEPES, 20 mM D-glucose, and 1 mM MgCl_2_ [pH 7.1]) at RT in the presence or absence of CIP. A 96-well microtiter plate was filled with around 200 µL of parasite suspension per well.

The dye-loaded cells fluoresced at 515 nm after being stimulated at 340 and 380 nm. Parasites loaded with SBFI were suspended in calibration buffers containing [Na^+^] values ranging from 0 to 140 mM (pH 7.1) to establish calibration curves (Diarra et al., 2001).

The cytosolic pH of wild-type and mutant strains was measured using the pH-sensitive fluorescent dye BCECF [2’,7’-bis-(2-carboxyethyl)-5-(and-6)- carboxyfluorescein] (Biotium; product 51011). The BCECF was added to saponin-isolated parasites by suspension (1×10^8^ parasites/mL) and incubated for 20 minutes at 37 °C in RPMI-1640 culture medium (Gibco, USA) (Saliba and Kirk, 1999). Thereafter, the parasites loaded with dye were rinsed thrice (12,000 × g, 1 min) in RPMI-1640 culture medium, then resuspended in physiological saline (as previously mentioned) with or without various concentrations of CIP. A 96-well microtiter plate was filled with around 200 µL of parasite suspension per well. The dye-loaded cells fluoresced at 520 nm after being stimulated at 440 and 490 nm. A pH calibration was carried out for each experiment (Mohring et al., 2022).

### Measurements of membrane ATPase activity

Membranes from isolated *B. gibsoni* were prepared by lysing the parasites in ice-cold deionized water containing 7 × Protease Inhibitor Cocktail Tablets (Roche, Germany). The membrane preparation was then washed three times with ice-cold deionized water, with protease inhibitors included in the first two washes (Qiu et al., 2022). Protein concentrations in the membrane samples were determined using a Bradford assay. The production of inorganic phosphate (Pi) from ATP hydrolysis was measured using the Malachite Green Phosphate Assay (BioAssay Systems, USA). Membrane preparations were diluted in either a high Na^+^ solution (final reaction conditions: 150 mM NaCl, 20 mM KCl, 2 mM MgCl_2_, 50 mM Tris, pH 7.2) or a Na^+^-free solution (final reaction conditions: 150 mM choline chloride, 20 mM KCl, 2 mM MgCl_2_, 50 mM Tris, pH 7.2) to achieve a final protein concentration of 40 μg/mL. CIP was added at the concentrations specified in the respective figure legends. Reactions were conducted according to the manufacturer’s protocol provided with the assay kit.

### Multiple ATP4 sequence alignment and molecular docking

*B. gibsoni* ATP4 (GenBank: KAK1443404.1), *B. bovis* ATP4 (PiroplasmaDB: BBOV_IV010020), *B. microti* ATP4 (GenBank: BMR1_03g01005), *T. gondii* ATP4 (GenBank: XP_018635122.1), and *Homo sapiens* ATP4 (GenBank: NM_000704.3) sequences were obtained by a homology search using *P. falciparum* ATP4 (GenBank: PF3D7_1211900). Sequence alignment was analyzed using MUSCLE in Jalview v2.11.3.2 software and BLAST (http://www.ncbi.nlm.nih.gov/BLAST/).

AlphaFold was used to predict the structure of BgATP4^WT^ (Jumper et al., 2021). Using the GROMACS 2021 Molecular Dynamics package, the energy minimization was carried out following the model’s generation (Lindahl et al., 2021). The mutations were produced by using Charmm-GUI PDB reader (Jo et al., 2014). PyMOL (version 2.0 Schrödinger, LLC) was used to confirm the position of the mutation site (center: -20.367, 10.904, 9.435; size: 30×30×30) (Trott and Olson, 2010). The ligand molecules CIP was downloaded from PubChem (CID 44469321 (https://pubchem.ncbi.nlm.nih.gov/compound/44469321). The ligand was used to dock with Gnina (Eberhardt et al., 2021). The affinity score and binding pose were chosen only from the highest CNN (convolutional neural network) score results from docking simulations. The models were visualized using the PyMOL Molecular Graphic System and Discovery Studio.

### Cross-resistance to TQ and ATO in mutant parasites

The efficacy of TQ and ATO against BgATP4^WT^, BgATP4^L921V^, and BgATP4^L921I^ was evaluated using a fluorescence assay, as described above. The IC_50_ values were calculated from the fluorescence values using non-linear regression analysis (curve fitting) in GraphPad Prism 9 (GraphPad Software Inc., USA).

### Efficacy of CIP plus TQ combination therapy in SCID mice infected with *B. microti*

Twenty 6-week-old female immunocompromised mice (C.B-17/IcrJcl-Prkdcscid strain, CLEA Japan, Inc.) were intraperitoneally injected with 1.0 × 10^7^ of *B. microti*-infected blood cells (Tuvshintulga et al., 2022). When the average parasitemia across all mice reached 1% (at 4 DPI), the drug treatments were initiated. The CIP group (n = 5) was orally administered 20 mg/kg CIP for seven days. The TQ group (n = 5) was orally administered a single dose of 20 mg/kg TQ. The CIP plus TQ group (n = 5) received the combination treatment, following the dosage and administration methods described above. The vehicle group (n = 5) orally received 0.2 mL of sesame oil as the control. Thin blood smears were prepared every other day and examined under a light microscope to determine parasitemia levels.

### qPCR analysis

To further verify the presence or absence of *B. microti* DNA in treated SCID mice, real-time quantitative PCR analysis was performed as described previously (Vydyam et al., 2024). Briefly, genomic DNA was extracted from blood samples (200 μl) collected at 90 DPI and subjected to qPCR analysis using SYBR Green I, targeting a highly conserved region of *Babesia* mitochondrial genome (mtDNA). Primers used were: Bmic-F 5′-TTGCGATAGTAATAGATTTACTGC-3′ and B-lsu-R2 5′-TCTTAACCCAACTCACGTACCA-3′ (Qurollo et al., 2017). The reaction mixture consisted of: 1 × Advanced Universal SYBR Green Super Mix (2 ×) (1725270; Bio-Rad); 0.5 μM of each primer, and 100 ng of genomic DNA.

### Detection of *B. microti* ATP4 gene mutations in relapsed infected SCID mice

The genomic DNA of the mutant parasites was extracted and sequenced (Ji et al., 2022a). The primer sets used for sequencing are listed in Table supplement 2. The single nucleotide variants were confirmed by pairwise alignment to the *Bm*ATP4^WT^ sequence (GenBank: BMR1_03g01005).

### Statistical analysis

Data analysis, namely one-way analysis of variance (ANOVA) and two-tailed unpaired t-tests, was performed using GraphPad Prism (La Jolla, CA, USA) version 9. A *P* value of <0.05 was considered a statistically significant result.

## Supporting information

Hightlighted version

## Acknowledgments

This work was supported by a Grant-in-Aid for Scientific Research (22H0250906), the JSPS Core-to-Core program, both from the Ministry of Education, Culture, Sports, Science, and Technology of Japan, and a grant from the Strategic International Collaborative Research Project (JPJ008837) promoted by the Ministry of Agriculture, Forestry, and Fisheries of Japan, and International Science and Technology Cooperation Project of Hubei Province (2024EHA018).

## Author contributions

Hang Li, Investigation, Data curation, Formal analysis, Writing – original draft, Writing – review & editing; Shengwei Ji, Methodology, Investigation, Software, Validation; Nanang R. Ariefta, Methodology, Software, Formal analysis; Eloiza May S. Galon, Visualization, Writing – review & editing; Shimaa AES El-Sayed, Methodology, Visualization; Thom Do, Methodology, Investigation; Lijun Jia, Data curation, Visualization; Miako Sakaguchi, Methodology, Visualization; Masahito Asada, Investigation, Visualization; Yoshifumi Nishikawa, Methodology, Resources; Xin Qin, Methodology, Data curation; Mingming Liu, Conceptualization, Methodology, Investigation, Resources, Data curation, Funding acquisition, Supervision, Writing – review & editing; Xuenan Xuan, Conceptualization, Methodology, Validation, Formal analysis, Supervision, Project administration, Writing – review & editing.

## Ethics

All animal experiments were performed according to the Guide for the Care and Use of Laboratory Animals of Obihiro University of Agriculture and Veterinary Medicine, which was also approved by the Committee on the Ethics of Animal Experiments at the Obihiro University of Agriculture and Veterinary Medicine, Japan (permit numbers: animal experiment, 22-145 and 23-132; DNA experiment, 2207 and 2208; pathogen, 202308 and 202306).

## Supplementary Materials

**Figure supplement 1.**
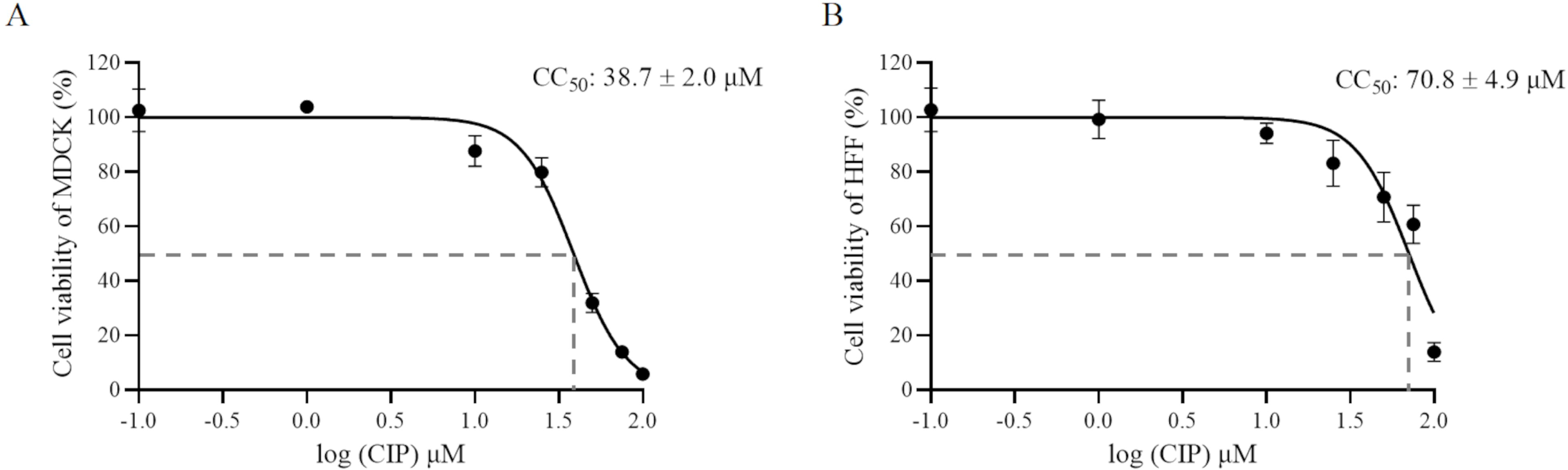
Cytotoxicity assay of CIP on the Madin-Darby canine kidney (MDCK) cells and human foreskin fibroblasts (HFF). The MTP-500 microplate reader was utilized to detect the absorbance at 450 nm. The results are displayed as the mean ± standard deviation of three separate tests. CC_50_: the 50% cytotoxic concentration.

**Figure supplement 2.**
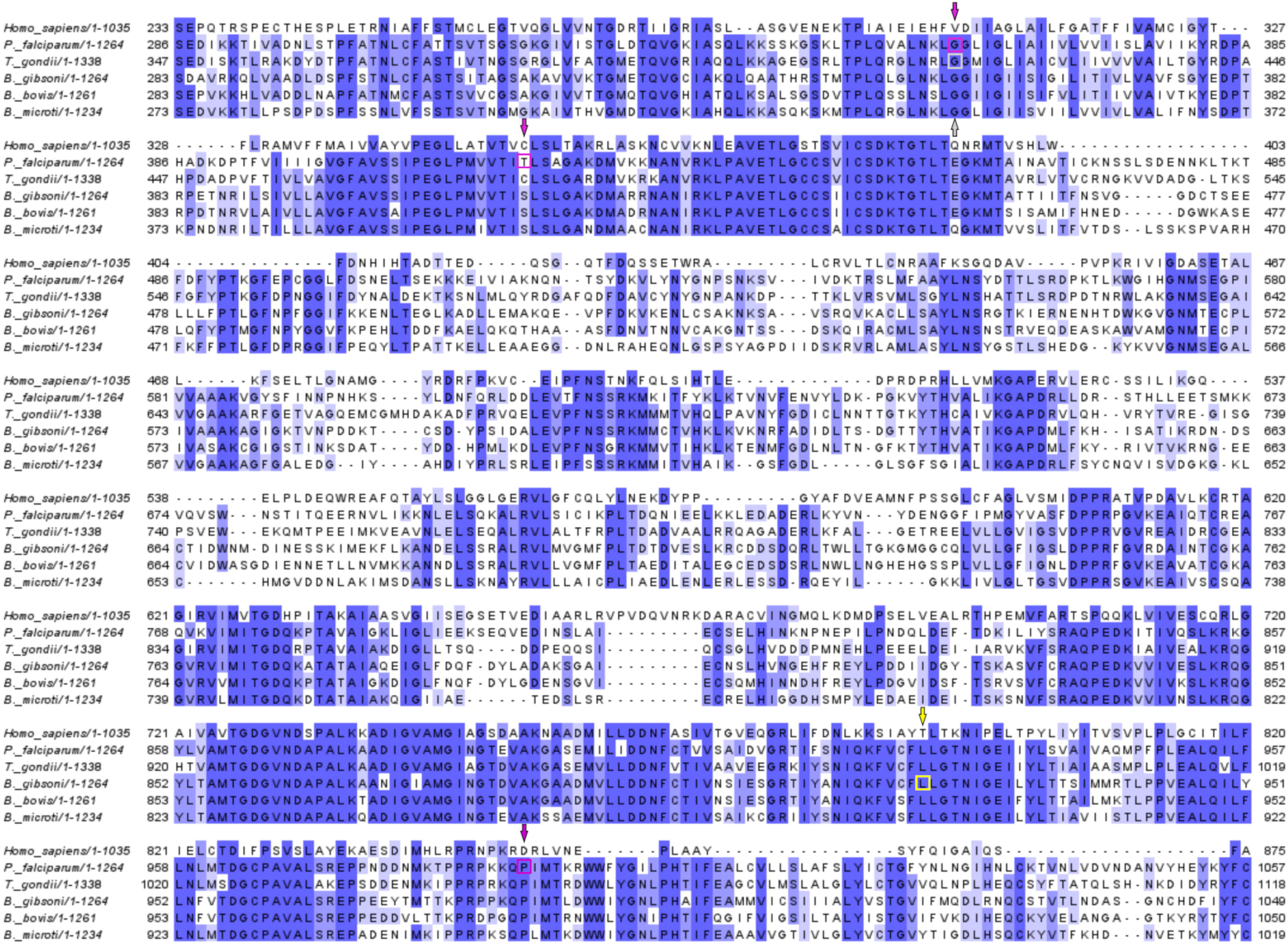
Multiple sequence alignment of ATP4 in different species. A yellow square and arrow denotes the BgATP4 mutation site discovered in this investigation; purple squares and arrows represent sites linked to *P. falciparum* CIP resistance, and a gray square and arrow represent sites associated with *T. gondii*.

**Figure supplement 3.**
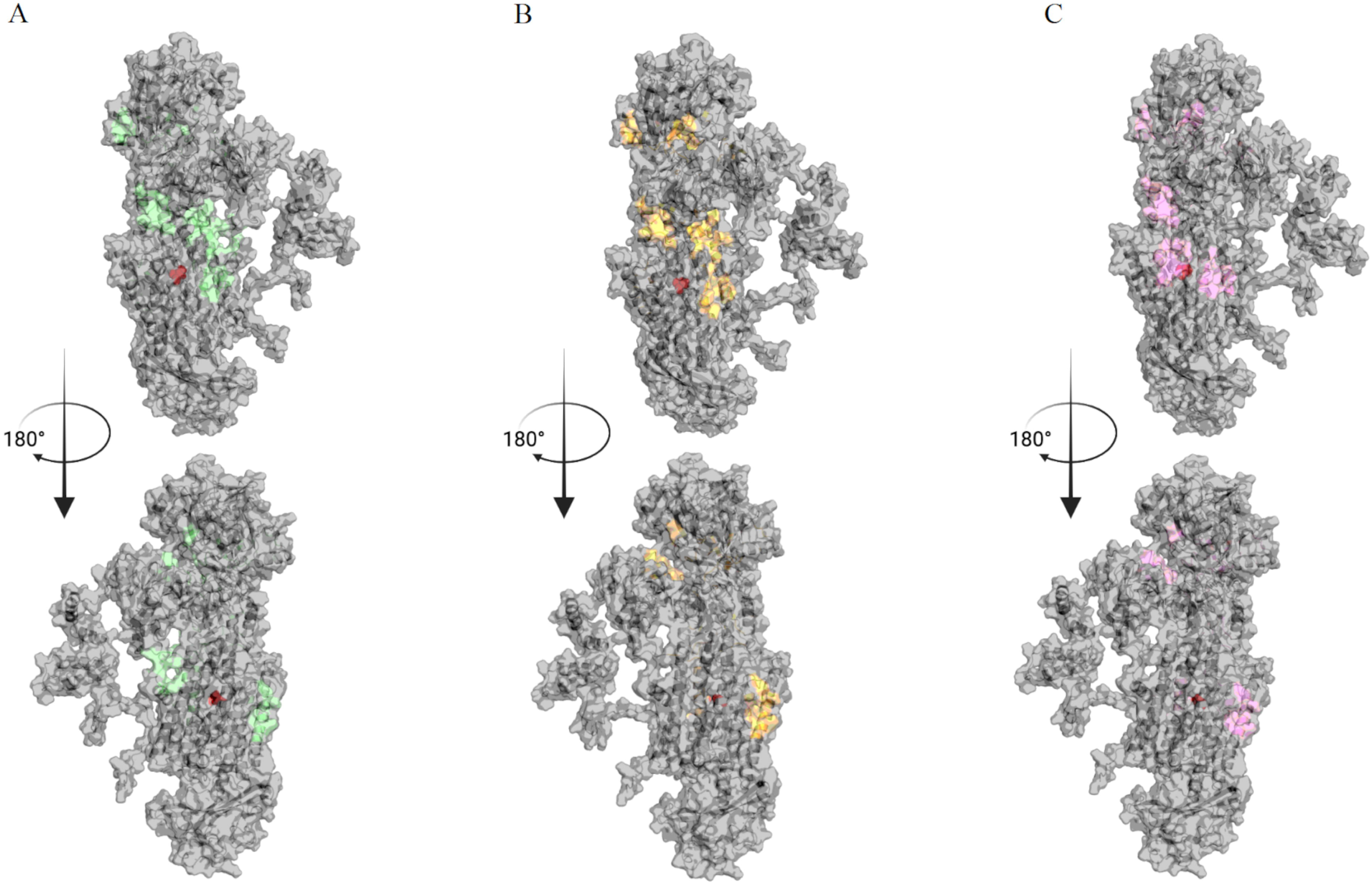
Binding sites for CIP found by Gnina search across the entire surface of the protein. (**A**-**C**) The binding space of WT, L921V, and L921I mutants in BgATP4 are labeled in green, yellow, and pink, respectively.

**Figure supplement 4.**
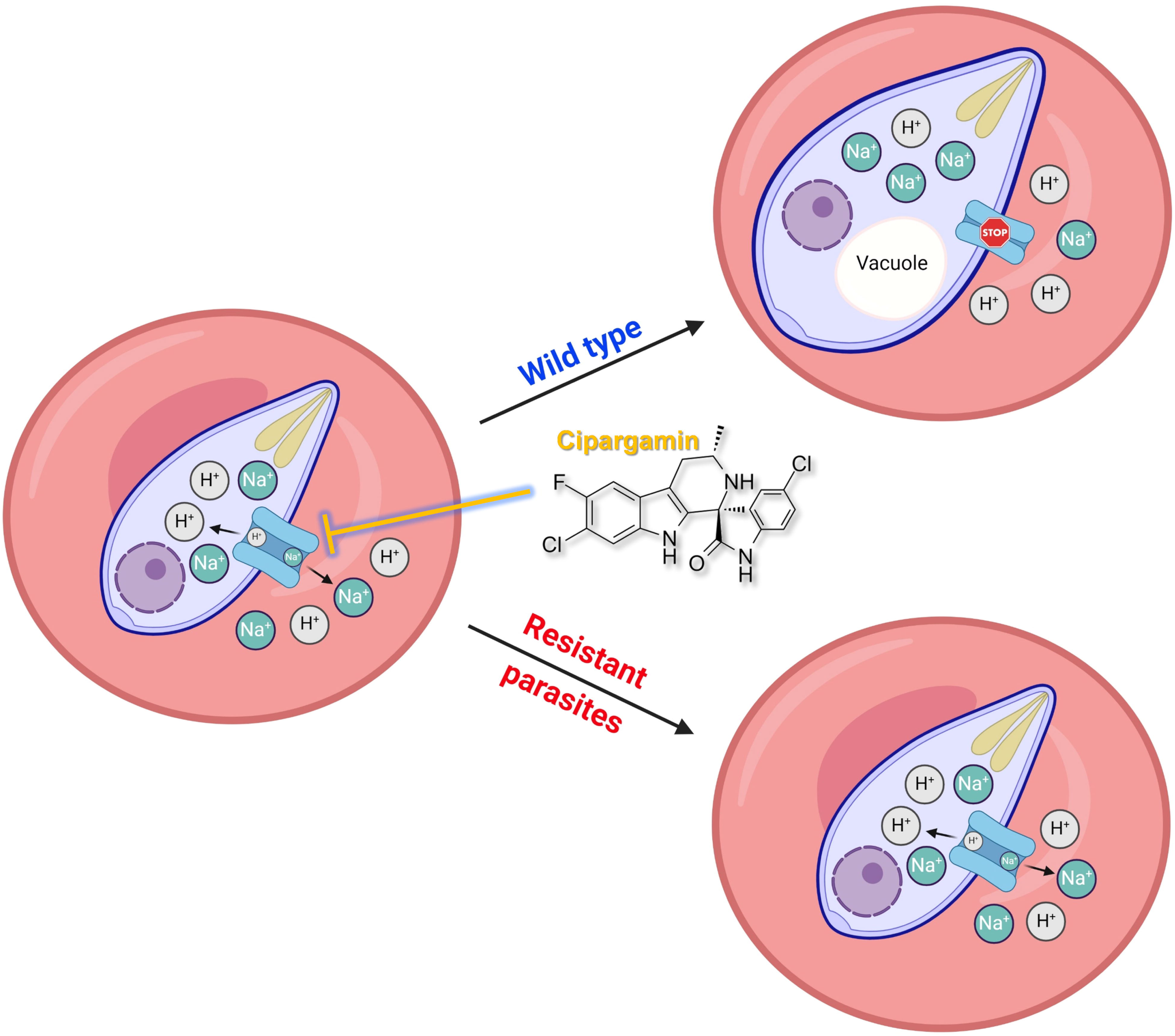
Proposed mechanism of inhibition of CIP on wild-type and mutant parasite-infected erythrocytes. CIP disrupts the BgATP4 function of wild-type parasites, which causes a net influx of Na^+^ and efflux of H^+^ from the parasite. The osmotic load imposed on the influx of Na^+^ further brings about parasite swelling and internal alkalinization, which are the main factors in *Babesia* death. Mutations in ATP4 minimize the susceptibility to ATP4 inhibitors by recovering H^+^ and Na^+^ balance, as indicated by the dotted arrows.

**Table supplement 1.**
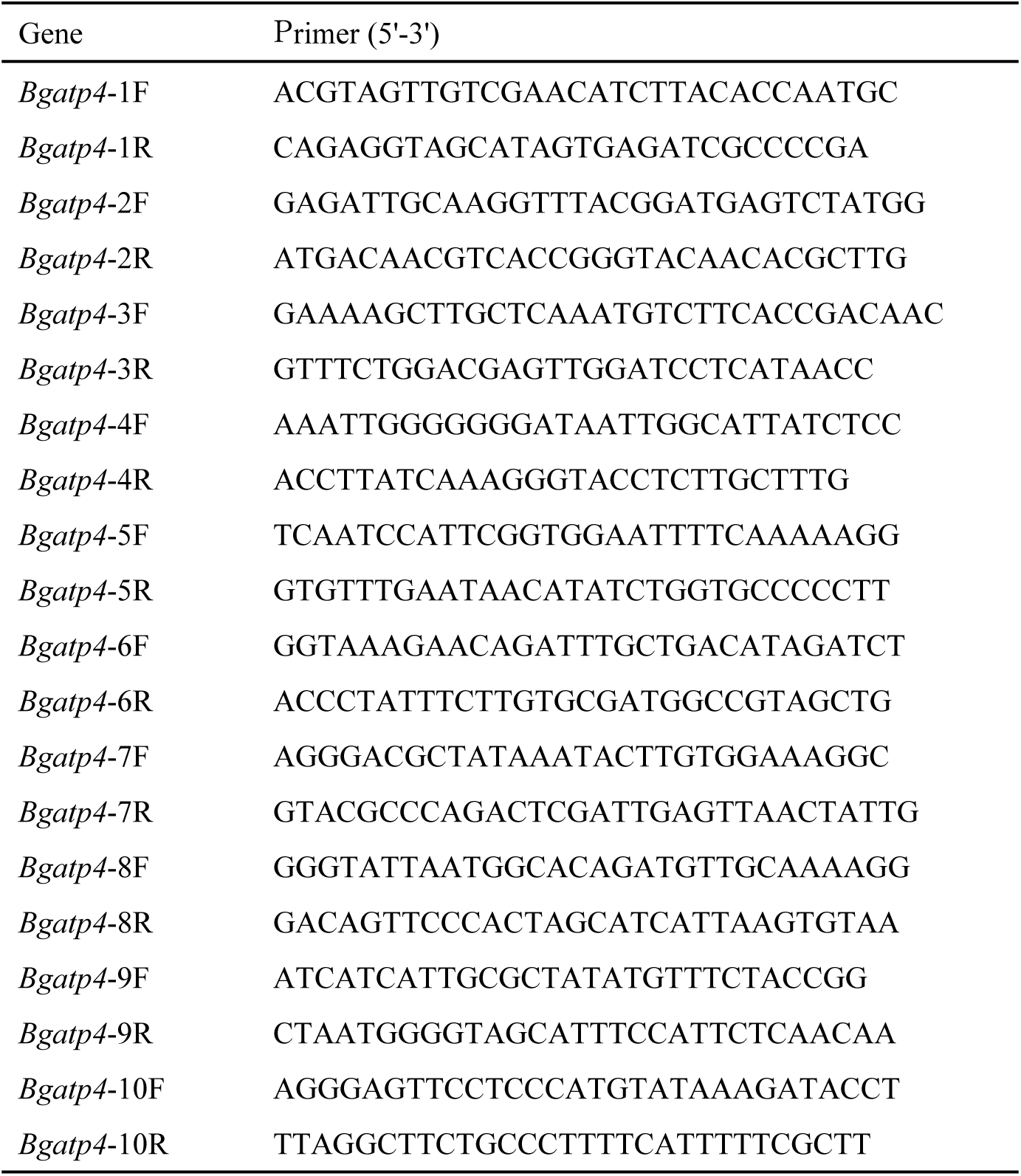
Primer sets of *B. gibsoni* ATP4.

**Table supplement 2.**
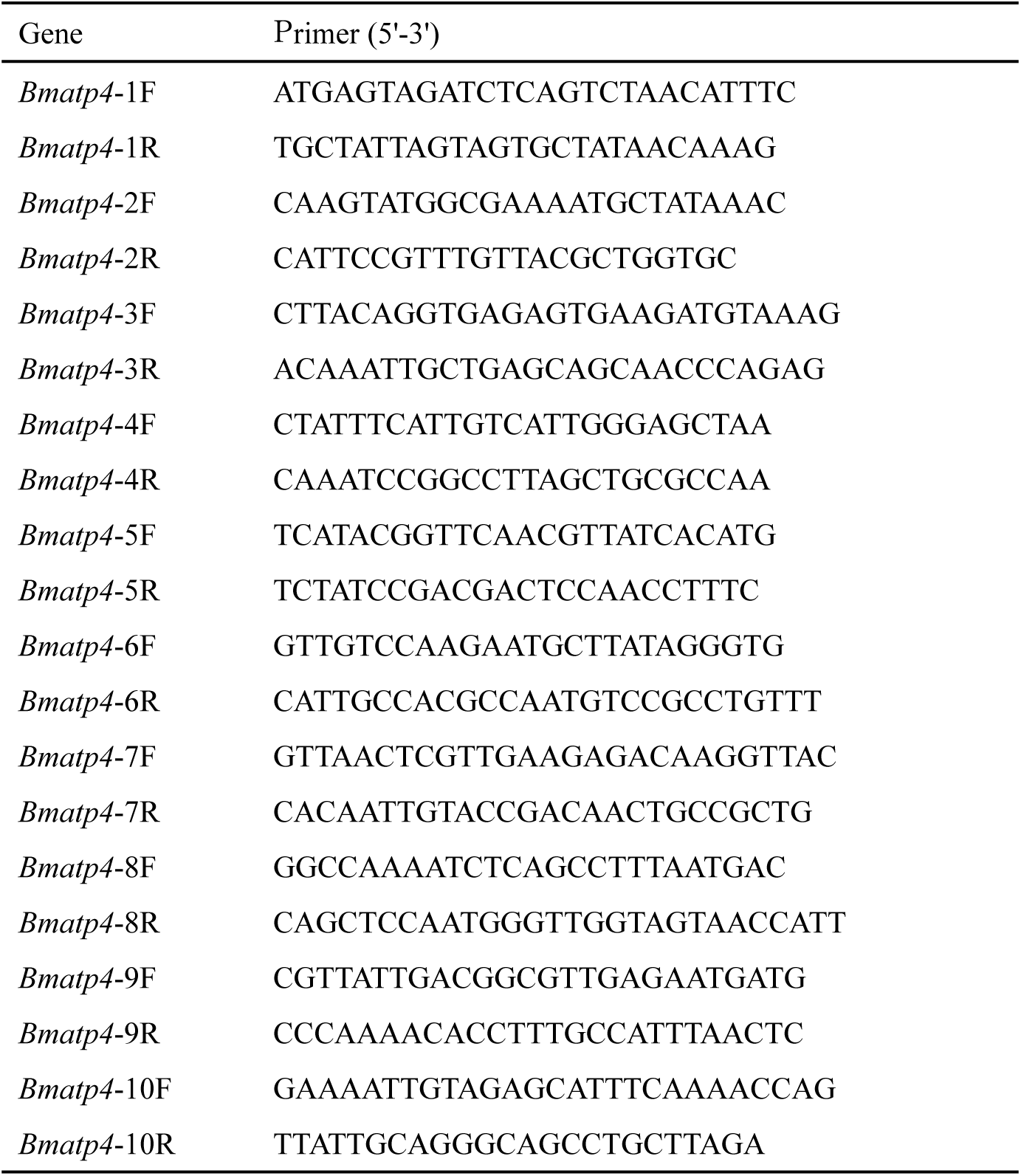
Primer sets of *B. microti* ATP4.

**Table supplement 3.**
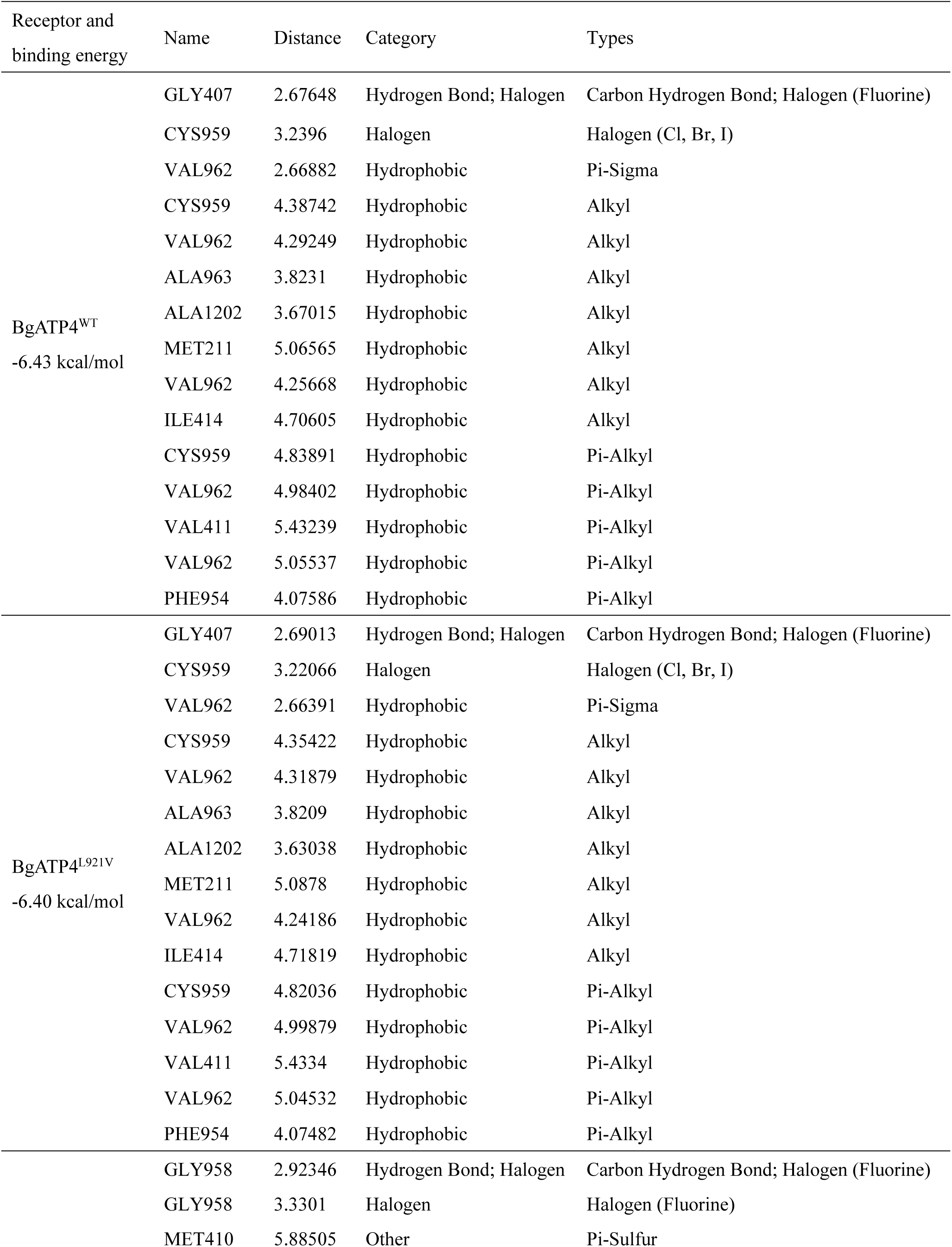
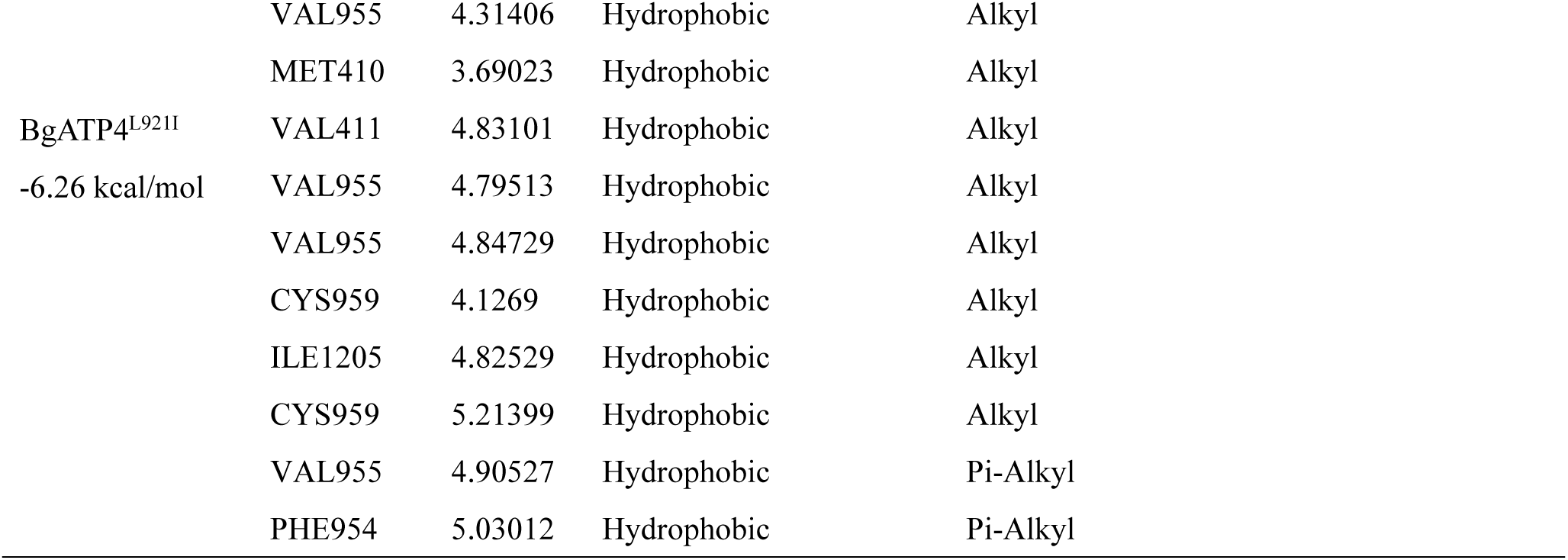
Interactions of CIP from docking simulations.

