## Supplementary material for "Efficacy and mechanism of action of cipargamin as an antibabesial drug candidate": Hightlighted version

**Address:** Longzhong Road No.296, Xiangyang, China; Nishi 11 Minami 39 Chome, Obihiro, Hokkaido

**Telephone number:** +8618904416800; +8109079036369

**Competing interest:** The authors have declared that no conflict of interest exists.

**Selection of CIP-resistant *B. gibsoni in vitro***

Selections were initiated by exposing six independent flasks, each containing 10 μL ($5\times10^6$) *B. gibsoni* iRBCs mixed with 40 μL ($4\times10^8$) RBCs into a 450 μL culture medium, which contained increasing concentrations of CIP: 5, 10, 20, 30 to 694 nM ($10\times IC_{50}$). The medium containing CIP was replaced daily until parasites treated with $10\times IC_{50}$ CIP reached multiplication rates that were approximately comparable to those of the untreated controls (Hwang et al., 2010). Then, to evaluate the decreased sensitivity to CIP, $IC_{50}$ values of resistant strains were determined by nonlinear regression using the GraphPad Prism software.

### Declaration of competing interest

The authors declare that they have no known competing financial interests or personal relationships that could have appeared to influence the work reported in this paper.

### Author contributions

Hang Li, Investigation, Data curation, Formal analysis, Writing – original draft, Writing – review & editing; Shengwei Ji, Methodology, Investigation, Software, Validation; Nanang R. Ariefta, Methodology, Software, Formal analysis; Eloiza May S. Galon, Visualization, Writing – review & editing; Shimaa AES El-Sayed, Methodology, Visualization; Thom Do, Methodology, Investigation; Lijun Jia, Data curation, Visualization; Miako Sakaguchi, Methodology, Visualization; Masahito Asada, Investigation, Visualization; Yoshifumi

Nishikawa, Methodology, Resources; Xin Qin, Methodology, Data curation; Mingming Liu, Conceptualization, Methodology, Investigation, Resources, Data curation, Funding acquisition, Supervision, Writing – review & editing; Xuenan Xuan, Conceptualization, Methodology, Validation, Formal analysis, Supervision, Project administration, Writing – review & editing.

### Ethics

All animal experiments were performed according to the Guide for the Care and Use of Laboratory Animals of Obihiro University of Agriculture and Veterinary Medicine, which was also approved by the Committee on the Ethics of Animal Experiments at the Obihiro University of Agriculture and Veterinary Medicine, Japan (permit numbers: animal experiment, 22-145 and 23-132; DNA experiment, 2207 and 2208; pathogen, 202308 and 202306).

ClinicalTrials.gov [Internet]. Bethesda (MD): National Library of Medicine (US). Identifier

NCT04675931, To evaluate efficacy, safety, tolerability and PK of intravenous cipargamin in participants with severe *Plasmodium falciparum* malaria; 2024 Feb 29 [cited 2024 Apr 9]. Available from: https://clinicaltrials.gov/study/NCT04675931

Das S, Bhatanagar S, Morrisey JM, Daly TM, Burns JM, Coppens I, Vaidya AB. 2016. $Na^+$ influx induced by new antimalarials causes rapid alterations in the cholesterol content and morphology of *Plasmodium falciparum*. *PLoS Pathog* **12**:e1005647. doi:10.1371/journal.ppat.1005647

Dennis ASM, Lehane AM, Ridgway MC, Holleran JP, Kirk K. 2018. Cell swelling induced by the antimalarial KAE609 (cipargamin) and other *Pf*ATP4-associated antimalarials. *Antimicrob Agents Chemother* **62**:e00087-18. doi:10.1128/AAC.00087-18

Diarra A, Sheldon C, Church J. 2001. In situ calibration and $[H^+]$ sensitivity of the fluorescent $Na^+$ indicator SBFI. *American Journal of Physiology-Cell Physiology* **280**:C1623–C1633. doi:10.1152/ajpcell.2001.280.6.C1623

Eberhardt J, Santos-Martins D, Tillack AF, Forli S. 2021. AutoDock Vina 1.2.0: new docking methods, expanded force field, and python bindings. *J Chem Inf Model* **61**:3891–3898. doi:10.1021/acs.jcim.1c00203

Gray JS, Ogden NH. 2021. Ticks, human babesiosis and climate change. *Pathogens* **10**:1430. doi:10.3390/pathogens10111430

Guswanto A, Sivakumar T, Rizk MA, Elsayed SAE, Youssef MA, ElSaid EES, Yokoyama N, Igarashi I. 2014. Evaluation of a fluorescence-based method for antibabesial drug screening. *Antimicrob Agents Chemother* **58**:4713–4717. doi:10.1128/AAC.00022-14

Hidayati AR, Melinda, Ilmi H, Sakura T, Sakaguchi M, Ohmori J, Hartuti ED, Tumewu L, Inaoka DK, Tanjung M, Yoshida E, Tokumasu F, Kita K, Mori M, Dobashi K, Nozaki T, Syafruddin D, Hafid AF, Waluyo D, Widyawaruyanti A. 2023. Effect of geranylated dihydrochalcone from *Artocarpus altilis* leaves extract on *Plasmodium falciparum* ultrastructural changes and mitochondrial malate: Quinone oxidoreductase. *International Journal for Parasitology: Drugs and Drug Resistance* **21**:40–50. doi:10.1016/j.ijpddr.2022.12.001

Holbrook NR, Klontz EH, Adams GC, Schnittman SR, Issa NC, Bond SA, Branda JA, Lemieux JE. 2023. *Babesia microti* variant with multiple resistance mutations detected in an immunocompromised patient receiving atovaquone prophylaxi. *Open Forum Infectious Diseases* **10**:ofad097. doi:10.1093/ofid/ofad097

Hwang S-J, Yamasaki M, Nakamura K, Sasaki N, Murakami M, Wickramasekara Rajapakshage BK, Ohta H, Maede Y, Takiguchi M. 2010. Development and characterization of a strain of *Babesia gibsoni* resistant to diminazene aceturate *in vitro*. *J Vet Med Sci* **72**:765–771. doi:10.1292/jvms.09-0535

Jalovecka M, Sojka D, Ascencio M, Schnittger L. 2019. *Babesia* life cycle-when phylogeny meets biology. *Trends in Parasitology* **35**:356–368. doi:10.1016/j.pt.2019.01.007

Jeon SA, Park JL, Park S-J, Kim JH, Goh S-H, Han J-Y, Kim S-Y. 2021. Comparison between MGI and Illumina sequencing platforms for whole genome sequencing. *Genes Genom* **43**:713–724. doi:10.1007/s13258-021-01096-x

Ji S, Galon EM, Rizk MA, Yi Y, Zafar I, Li H, Ma Z, Iguchi A, Asada M, Liu M, Xuan X. 2022a. Efficacy of the antimalarial MMV390048 against *Babesia* infection reveals phosphatidylinositol 4-kinase as a druggable target for babesiosis. *Antimicrob Agents Chemother* **66**:e00574-22. doi:10.1128/aac.00574-22

Ji S, Liu M, Galon EM, Rizk MA, Tuvshintulga B, Li J, Zafar I, Hasegawa Y, Iguchi A, Yokoyama N, Xuan X. 2022b. Inhibitory effect of naphthoquine phosphate on *Babesia gibsoni in vitro* and *Babesia rodhaini in vivo*. *Parasites Vectors* **15**:10. doi:10.1186/s13071-021-05127-0

Jo S, Cheng X, Islam SM, Huang L, Rui H, Zhu A, Lee HS, Qi Y, Han W, Vanommeslaeghe K, MacKerell AD, Roux B, Im W. 2014. CHARMM-GUI PDB manipulator for advanced modeling and simulations of proteins containing nonstandard residues. *Adv Protein Chem Struct Biol* **96:** 235–265. doi:10.1016/bs.apcsb.2014.06.002

Jumper J, Evans R, Pritzel A, Green T, Figurnov M, Ronneberger O, Tunyasuvunakool K, Bates R, Žídek A, Potapenko A, Bridgland A, Meyer C, Kohl SAA, Ballard AJ, Cowie A, Romera-Paredes B, Nikolov S, Jain R, Adler J, Back T, et al. 2021. Highly accurate protein structure prediction with AlphaFold. *Nature* **596**:583–589. doi:10.1038/s41586-021-03819-2

Kletsova EA, Spitzer ED, Fries BC, Marcos LA. 2017. Babesiosis in Long Island: review of 62 cases focusing on treatment with azithromycin and atovaquone. *Ann Clin Microbiol Antimicrob* **16**:26. doi:10.1186/s12941-017-0198-9

Krause PJ. 2019. Human babesiosis. *International Journal for Parasitology* **49**:165–174. doi:10.1016/j.ijpara.2018.11.007

Krause PJ, Auwaerter PG, Bannuru RR, Branda JA, Falck-Ytter YT, Lantos PM, Lavergne V, Meissner HC, Osani MC, Rips JG, Sood SK, Vannier E, Vaysbrot EE, Wormser GP. 2021. Clinical practice guidelines by the infectious diseases society of America (IDSA): 2020 guideline on diagnosis and management of babesiosis. *Clinical Infectious Diseases* **72**:e49–e64. doi:10.1093/cid/ciaa1216

Krause PJ, Rogers R, Shah MK, Kang H, Parsonnet J, Kodama R, Vannier E. 2024. Tafenoquine for relapsing babesiosis: a case series. *Clinical Infectious Diseases* **79**:130–137. doi:10.1093/cid/ciae238

Krishna S, Woodrow C, Webb R, Penny J, Takeyasu K, Kimura M, East JM. 2001. Expression and functional characterization of a *Plasmodium falciparum* $Ca2^{+}$-ATPase (*Pf*ATP4) belonging to a subclass unique to apicomplexan organisms. *Journal of Biological Chemistry* **276**:10782–10787. doi:10.1074/jbc.M010554200

Lee AH, Fidock DA. 2016. Evidence of a mild mutator phenotype in cambodian *Plasmodium* *falciparum* malaria parasites. *PLoS ONE* **11**:e0154166. doi:10.1371/journal.pone.0154166

Lehane AM, Dennis ASM, Bray KO, Li D, Rajendran E, McCoy JM, McArthur HM, Winterberg M, Rahimi F, Tonkin CJ, Kirk K, Van Dooren GG. 2019. Characterization of the ATP4 ion pump in *Toxoplasma gondii*. *Journal of Biological Chemistry* **294**:5720–5734. doi:10.1074/jbc.RA118.006706

Lemieux JE, Tran AD, Freimark L, Schaffner SF, Goethert H, Andersen KG, Bazner S, Li A, McGrath G, Sloan L, Vannier E, Milner D, Pritt B, Rosenberg E, Telford S, Bailey JA, Sabeti PC. 2016. A global map of genetic diversity in *Babesia microti* reveals strong population structure and identifies variants associated with clinical relapse. *Nat Microbiol* **1**:16079. doi:10.1038/nmicrobiol.2016.79

Li H, Galon EM, Ji S, Zafar I, Ma Z, Do T, Amer MM, Ma Y, Liu M, Xuan X. 2023. *In vitro* screening of compounds from the Food and Drug Administration-approved library identifies anti-*Babesia gibsoni* activity of idarubicin hydrochloride and vorinostat. *Parasitology International* **96**:102774. doi:10.1016/j.parint.2023.102774

Lindahl, Abraham, Hess, Spoel VD. 2021. GROMACS 2021 Manual. doi:10.5281/ZENODO.4457591

Liu M, Adjou Moumouni PF, Cao S, Asada M, Wang G, Gao Y, Guo H, Li J, Vudriko P, Efstratiou A, Ringo AE, Lee S-H, Hakimi H, Masatani T, Sunaga F, Kawazu S, Yamagishi J, Jia L, Inoue N, Xuan X. 2018. Identification and characterization of interchangeable cross-species functional promoters between *Babesia gibsoni* and *Babesia bovis*. *Ticks and Tick-borne Diseases* **9**:330–333. doi:10.1016/j.ttbdis.2017.11.008

Liu M, Ji S, Kondoh D, Galon EM, Li J, Tomihari M, Yanagawa M, Tagawa M, Adachi M, Asada M, Igarashi I, Iguchi A, Xuan X. 2021. Tafenoquine is a promising drug candidate for the treatment of babesiosis. *Antimicrob Agents Chemother* **65**:e00204-21. doi:10.1128/AAC.00204-21

Marcos LA, Leung A, Kirkman L, Wormser GP. 2022. Use of tafenoquine to treat a patient with relapsing babesiosis with clinical and molecular evidence of resistance to azithromycin and atovaquone. *IDCases* **27**:e01460. doi:10.1016/j.idcr.2022.e01460

Matsuu A, Koshida Y, Kawahara M, Inoue K, Ikadai H, Hikasa Y, Okano S, Higuchi S. 2004. Efficacy of atovaquone against *Babesia gibsoni in vivo* and *in vitro*. *Veterinary Parasitology* **124**:9-18. doi:10.1016/j.vetpar.2004.07.005

Mohring F, Van Schalkwyk DA, Henrici RC, Blasco B, Leroy D, Sutherland CJ, Moon RW. 2022. Cation ATPase (ATP4) orthologue replacement in the malaria parasite *Plasmodium knowlesi* reveals species-specific responses to ATP4-targeting drugs. *mBio* **13**:e01178-22. doi:10.1128/mbio.01178-22

Mordue DG, Wormser GP. 2019. Could the drug tafenoquine revolutionize treatment of *Babesia microti* infection? *J Infect Dis* **220**:442-447. doi: 10.1093/infdis/jiz119

Ndayisaba G, Yeka A, Asante KP, Grobusch MP, Karita E, Mugerwa H, Asiimwe S, Oduro A,

Fofana B, Doumbia S, Jain JP, Barsainya S, Kullak-Ublick GA, Su G, Schmitt EK, Csermak K, Gandhi P, Hughes D. 2021. Hepatic safety and tolerability of cipargamin (KAE609), in adult patients with *Plasmodium falciparum* malaria: a randomized, phase II, controlled, dose-escalation trial in sub-Saharan Africa. *Malar J* **20**:478. doi:10.1186/s12936-021-04009-1

Qiu D, Pei JV, Rosling JEO, Thathy V, Li D, Xue Y, Tanner JD, Penington JS, Aw YTV, Aw JYH, Xu G, Tripathi AK, Gnadig NF, Yeo T, Fairhurst KJ, Stokes BH, Murithi JM, Kümpornsin K, Hasemer H, Dennis ASM, et al. 2022. A G358S mutation in the *Plasmodium falciparum* $Na^+$ pump *Pf*ATP4 confers clinically-relevant resistance to cipargamin. *Nat Commun* **13**:5746. doi:10.1038/s41467-022-33403-9

Qurollo BA, Archer NR, Schreeg ME, Marr HS, Birkenheuer AJ, Haney KN, Thomas BS, Breitschwerdt EB. 2017. Improved molecular detection of *Babesia* infections in animals using a novel quantitative real-time PCR diagnostic assay targeting mitochondrial DNA. *Parasites Vectors* **10**:128. doi:10.1186/s13071-017-2064-1

Rogers R, Krause PJ, Norris AM, Ting MH, Nagami EH, Cilley B, Vannier E. 2023. Broad antimicrobial resistance in a case of relapsing babesiosis successfully treated with tafenoquine. *Clinical Infectious Diseases* **76**:741–744. doi:10.1093/cid/ciac473

Rosenblatt J, Leung A, Baneman E, Fuller R, Taimur S, Paniz-Mondolfi AE, Malone AK, Kirkman L, Jacobs SE. 2021. Relapsed *Babesia microti* infection following allogeneic hematopoietic cell transplantation in a patient with B-cell acute lymphoblastic leukemia: case report and review of the literature. *Open Forum Infectious Diseases* **8**:ofab323. doi:10.1093/ofid/ofab323

Rosling JEO, Ridgway MC, Summers RL, Kirk K, Lehane AM. 2018. Biochemical characterization and chemical inhibition of *Pf*ATP4-associated $Na^+$-ATPase activity in *Plasmodium falciparum* membranes. *Journal of Biological Chemistry* **293**:13327–13337. doi:10.1074/jbc.RA118.003640

Rottmann M, McNamara C, Yeung BKS, Lee MCS, Zou B, Russell B, Seitz P, Plouffe DM, Dharia NV, Tan J, Cohen SB, Spencer KR, González-Páez GE, Lakshminarayana SB, Goh A, Suwanarusk R, Jegla T, Schmitt EK, Beck H-P, Brun R, et al. 2010. Spiroindolones, a potent compound class for the treatment of malaria. *Science* **329**:1175–1180. doi:10.1126/science.1193225

Saliba KJ, Kirk K. 1999. pH regulation in the intracellular malaria parasite, *Plasmodium falciparum*. *Journal of Biological Chemistry* **274**:33213–33219. doi:10.1074/jbc.274.47.33213

Schmitt EK, Ndayisaba G, Yeka A, Asante KP, Grobusch MP, Karita E, Mugerwa H, Asiimwe S, Oduro A, Fofana B, Doumbia S, Su G, Csermak Renner K, Venishetty VK, Sayyed S, Straimer J, Demin I, Barsainya S, Boulton C, Gandhi P. 2022. Efficacy of cipargamin (KAE609) in a randomized, phase II dose-escalation study in adults in Sub-Saharan Africa with uncomplicated *Plasmodium falciparum* malaria. *Clinical Infectious Diseases* **74**:1831–1839. doi:10.1093/cid/ciab716

Simon MS, Westblade LF, Dziedziech A, Visone JE, Furman RR, Jenkins SG, Schuetz AN, Kirkman LA. 2017. Clinical and molecular evidence of atovaquone and azithromycin resistance in relapsed *Babesia microti* infection associated with rituximab and chronic lymphocytic leukemia. *Clinical Infectious Diseases* **65**:1222–1225. doi:10.1093/cid/cix477

Spillman NJ, Allen RJW, McNamara CW, Yeung BKS, Winzeler EA, Diagana TT, Kirk K. 2013. $Na^+$ regulation in the malaria parasite *Plasmodium falciparum* involves the cation ATPase *Pf*ATP4 and is a target of the spiroindolone antimalarials. *Cell Host & Microbe* **13**:227–237. doi:10.1016/j.chom.2012.12.006

Tayebwa DS, Tuvshintulga B, Guswanto A, Nugraha AB, Batiha GE-S, Gantuya S, Rizk MA, Vudriko P, Sivakumar T, Yokoyama N, Igarashi I. 2018. The effects of nitidine chloride and camptothecin on the growth of *Babesia* and *Theileria* parasites. *Ticks and Tick-borne Diseases* **9**:1192–1201. doi:10.1016/j.ttbdis.2018.04.019

Trott O, Olson AJ. 2010. AutoDock Vina: improving the speed and accuracy of docking with a new scoring function, efficient optimization, and multithreading. *J Comput Chem* **31**:455–461. doi:10.1002/jcc.21334

Tuvshintulga B, Sivakumar T, Nugraha AB, Ahedor B, Batmagnai E, Otgonsuren D, Liu M, Xuan X, Igarashi I, Yokoyama N. 2022. Combination of clofazimine and atovaquone as a potent therapeutic regimen for the radical cure of *Babesia microti* infection in immunocompromised hosts. *The Journal of Infectious Diseases* **225**:238–242. doi:10.1093/infdis/jiab537

Vannier E, Krause PJ. 2012. Human babesiosis. *N Engl J Med* **366**:2397–2407. doi:10.1056/NEJMra1202018

Vega CA, Buening GM, Green TJ, Carson CA. 1985. *In vitro* cultivation of *Babesia bigemina*. *Am J Vet Res* **46**:416–420.

Vydyam P, Pal AC, Renard I, Chand M, Kumari V, Gennaro JC, Mamoun CB. 2024. Tafenoquine-atovaquone combination achieves radical cure and confers sterile immunity in experimental models of human babesiosis. *The Journal of Infectious Diseases* **229**:161–172. doi:10.1093/infdis/jiad315

Zhang R, Suwanarusk R, Malleret B, Cooke BM, Nosten F, Lau Y-L, Dao M, Lim CT, Renia L, Tan KSW, Russell B. 2016. A basis for rapid clearance of circulating ring-stage malaria parasites by the spiroindolone KAE609. *J Infect Dis* **213**:100–104. doi:10.1093/infdis/jiv358

Zhou Y, Fomovska A, Muench S, Lai B-S, Mui E, McLeod R. 2014. Spiroindolone that inhibits *Pf*ATPase4 is a potent, cidal inhibitor of *Toxoplasma gondii* tachyzoites *in vitro* and *in vivo*. *Antimicrob Agents Chemother* **58**:1789–1792. doi:10.1128/AAC.02225-13

### Supplemental Tables

**Table supplement 1.** Primer sets of *B. gibsoni* ATP4

| Gene | Primer (5'-3') |
|---|---|
| *Bgatp4*-1F | ACGTAGTTGTCGAACATCTTACACCAATGC |
| *Bgatp4*-1R | CAGAGGTAGCATAGTGAGATCGCCCCGA |
| *Bgatp4*-2F | GAGATTGCAAGGTTTACGGATGAGTCTATGG |
| *Bgatp4*-2R | ATGACAACGTCACCGGGTACAACACGCTTG |
| *Bgatp4*-3F | GAAAAGCTTGCTCAAATGTCTTCACCGACAAC |
| *Bgatp4*-3R | GTTTCTGGACGAGTTGGATCCTCATAACC |
| *Bgatp4*-4F | AAATTGGGGGGGATAATTGGCATTATCTCC |
| *Bgatp4*-4R | ACCTTATCAAAGGGTACCTCTTGCTTTG |
| *Bgatp4*-5F | TCAATCCATTCGGTGGAATTTTCAAAAAGG |
| *Bgatp4*-5R | GTGTTTGAATAACATATCTGGTGCCCCCTT |
| *Bgatp4*-6F | GGTAAAGAACAGATTTGCTGACATAGATCT |
| *Bgatp4*-6R | ACCCTATTTCTTGTGCGATGGCCGTAGCTG |
| *Bgatp4*-7F | AGGGACGCTATAAATACTTGTGGAAAGGC |
| *Bgatp4*-7R | GTACGCCCAGACTCGATTGAGTTAACTATTG |
| *Bgatp4*-8F | GGGTATTAATGGCACAGATGTTGCAAAAGG |
| *Bgatp4*-8R | GACAGTTCCCACTAGCATCATTAAGTGTAA |
| *Bgatp4*-9F | ATCATCATTGCGCTATATGTTTCTACCGG |
| *Bgatp4*-9R | CTAATGGGGTAGCATTTCCATTCTCAACAA |
| *Bgatp4*-10F | AGGGAGTTCCTCCCATGTATAAAGATACCT |
| *Bgatp4*-10R | TTAGGCTTCTGCCCTTTTCATTTTTCGCTT |

**Table supplement 2.** Primer sets of *B. microti* ATP4

| Gene | Primer (5'-3') |
|---|---|
| *Bmatp4*-1F | ATGAGTAGATCTCAGTCTAACATTTC |
| *Bmatp4*-1R | TGCTATTAGTAGTGCTATAACAAAG |
| *Bmatp4*-2F | CAAGTATGGCGAAAATGCTATAAAC |
| *Bmatp4*-2R | CATTCCGTTTGTTACGCTGGTGC |
| *Bmatp4*-3F | CTTACAGGTGAGAGTGAAGATGTAAAG |
| *Bmatp4*-3R | ACAAATTGCTGAGCAGCAACCCAGAG |
| *Bmatp4*-4F | CTATTTCATTGTCATTGGGAGCTAA |
| *Bmatp4*-4R | CAAATCCGGCCTTAGCTGCGCCAA |
| *Bmatp4*-5F | TCATACGGTTCAACGTTATCACATG |
| *Bmatp4*-5R | TCTATCCGACGACTCCAACCTTTC |
| *Bmatp4*-6F | GTTGTCCAAGAATGCTTATAGGGTG |
| *Bmatp4*-6R | CATTGCCACGCCAATGTCCGCCTGTTT |
| *Bmatp4*-7F | GTTAACTCGTTGAAGAGACAAGGTTAC |
| *Bmatp4*-7R | CACAATTGTACCGACAACTGCCGCTG |
| *Bmatp4*-8F | GGCCAAAATCTCAGCCTTTAATGAC |
| *Bmatp4*-8R | CAGCTCCAATGGGTTGGTAGTAACCATT |
| *Bmatp4*-9F | CGTTATTGACGGCGTTGAGAATGATG |
| *Bmatp4*-9R | CCCAAAACACCTTTGCCATTTAACTC |
| *Bmatp4*-10F | GAAAATTGTAGAGCATTTCAAAACCAG |
| *Bmatp4*-10R | TTATTGCAGGGCAGCCTGCTTAGA |

**Table supplement 3.** Interactions of CIP from docking simulations

| Receptor and binding energy | Name | Distance | Category | Types |
|---|---|---|---|---|
| BgATP4$^{WT}$ -6.43 kcal/mol | GLY407 | 2.67648 | Hydrogen Bond; Halogen | Carbon Hydrogen Bond; Halogen (Fluorine) |
| | CYS959 | 3.2396 | Halogen | Halogen (Cl, Br, I) |
| | VAL962 | 2.66882 | Hydrophobic | Pi-Sigma |
| | CYS959 | 4.38742 | Hydrophobic | Alkyl |
| | VAL962 | 4.29249 | Hydrophobic | Alkyl |
| | ALA963 | 3.8231 | Hydrophobic | Alkyl |
| | ALA1202 | 3.67015 | Hydrophobic | Alkyl |
| | MET211 | 5.06565 | Hydrophobic | Alkyl |
| | VAL962 | 4.25668 | Hydrophobic | Alkyl |
| | ILE414 | 4.70605 | Hydrophobic | Alkyl |
| | CYS959 | 4.83891 | Hydrophobic | Pi-Alkyl |
| | VAL962 | 4.98402 | Hydrophobic | Pi-Alkyl |
| | VAL411 | 5.43239 | Hydrophobic | Pi-Alkyl |
| | VAL962 | 5.05537 | Hydrophobic | Pi-Alkyl |
| | PHE954 | 4.07586 | Hydrophobic | Pi-Alkyl |
| BgATP4$^{L921V}$ -6.40 kcal/mol | GLY407 | 2.69013 | Hydrogen Bond; Halogen | Carbon Hydrogen Bond; Halogen (Fluorine) |
| | CYS959 | 3.22066 | Halogen | Halogen (Cl, Br, I) |
| | VAL962 | 2.66391 | Hydrophobic | Pi-Sigma |
| | CYS959 | 4.35422 | Hydrophobic | Alkyl |
| | VAL962 | 4.31879 | Hydrophobic | Alkyl |
| | ALA963 | 3.8209 | Hydrophobic | Alkyl |
| | ALA1202 | 3.63038 | Hydrophobic | Alkyl |
| | MET211 | 5.0878 | Hydrophobic | Alkyl |
| | VAL962 | 4.24186 | Hydrophobic | Alkyl |
| | ILE414 | 4.71819 | Hydrophobic | Alkyl |
| | CYS959 | 4.82036 | Hydrophobic | Pi-Alkyl |
| | VAL962 | 4.99879 | Hydrophobic | Pi-Alkyl |
| | VAL411 | 5.4334 | Hydrophobic | Pi-Alkyl |
| | VAL962 | 5.04532 | Hydrophobic | Pi-Alkyl |
| | PHE954 | 4.07482 | Hydrophobic | Pi-Alkyl |
| | GLY958 | 2.92346 | Hydrogen Bond; Halogen | Carbon Hydrogen Bond; Halogen (Fluorine) |
| | GLY958 | 3.3301 | Halogen | Halogen (Fluorine) |
| | MET410 | 5.88505 | Other | Pi-Sulfur |

| | | | | |
|---|---|---|---|---|
| | VAL955 | 4.31406 | Hydrophobic | Alkyl |
| | MET410 | 3.69023 | Hydrophobic | Alkyl |
| BgATP4$^{L921I}$ | VAL411 | 4.83101 | Hydrophobic | Alkyl |
| -6.26 kcal/mol | VAL955 | 4.79513 | Hydrophobic | Alkyl |
| | VAL955 | 4.84729 | Hydrophobic | Alkyl |
| | CYS959 | 4.1269 | Hydrophobic | Alkyl |
| | ILE1205 | 4.82529 | Hydrophobic | Alkyl |
| | CYS959 | 5.21399 | Hydrophobic | Alkyl |
| | VAL955 | 4.90527 | Hydrophobic | Pi-Alkyl |
| | PHE954 | 5.03012 | Hydrophobic | Pi-Alkyl |
